# Diversity of an uncommon elastic hypersaline microbial mat along a small-scale transect

**DOI:** 10.1101/2021.03.04.433984

**Authors:** Laura Espinosa-Asuar, Camila Monroy-Guzmán, David Madrigal-Trejo, Marisol Navarro-Miranda, Jazmin Sánchez-Pérez, Jhoselinne Buenrostro-Muñoz, Juan Villar, Julián Felipe Cifuentes Camargo, Maria Kalambokidis, Diego A. Esquivel-Hernandez, Mariette Viladomat Jasso, Ana E. Escalante, Patricia Velez, Mario Figueroa, Anahí Martínez Cárdenas, Santiago Ramirez Barahona, Jaime Gasca-Pineda, Luis E. Eguiarte, Valeria Souza

## Abstract

We evaluated the microbial diversity and metabolome profile of an uncommon hypersaline elastic microbial mat from Cuatro Ciénegas Basin (CCB) in the Chihuahuan Desert of Coahuila, México. We collected ten samples on a small scale transect (1.5-meters) and described its microbial diversity through NGS-based ITS and 16S rDNA gene sequencing. A very low number of taxa comprised a considerable proportion of the mat and were shared across all sampling points, whereas the rare biosphere was more phylogenetically diverse (Faith’s Phylogenetic Diversity (FPD) index) and phylogenetically disperse (using a null model distribution of Phylogenetic Species Clustering (nmdPSC)) than the abundant (high read count) taxa for both analyzed libraries. We also found a distinctive metabolome profile for each sample and were able to tentatively annotate several classes of compounds with relevant biological properties.

## Introduction

Microbial mats are tightly interacting communities, usually self-sustaining, that are horizontally stratified with multilayered biofilms embedded in a matrix of exopolysaccharides and inorganic substances that bind cells together (Bolhuis et al., 2014; Prieto-Barajas et al., 2018; Stolz, 2000). These stratified structures, as well as stromatolites, are the oldest communities found in the fossil record, informing our understanding of ancient metabolic interactions (Gutiérrez-Preciado et al., 2018; Nutman et al., 2016).

Although modern microbial mats and stromatolites are geographically widespread (Gerdes, 2010), their distribution is restricted to extreme environmental conditions where multicellular algae and grazing organisms cannot grow (Bebout et al., 2002). One such site, with a high diversity of microbial mats and stromatolites, is the desert oasis of the Cuatro Ciénegas Basin (CCB), found in the Chihuahuan Desert of the state of Coahuila in northern México. This small site (150,000 km^2^) is extremely biodiverse – in particular for microbes (Souza, Eguiarte, et al., 2018) – and is considered of international importance by the RAMSAR convention (www.ramsar.org). CCB is characterized by an imbalanced stoichiometry (i.e., very low concentration of phosphorus) (Elser et al., 2005, 2006; Papineau, 2010; Planavsky et al., 2010), which appears to be a strong selective pressure driving microbial lineages to adapt locally (e.g. the reported *Bacillus coahuilensis* genes involved in phosphorous utilization efficiency (Alcaraz et al. 2008)). The uniqueness of microbial life in CCB -- deemed as an “Astrobiological Precambrian Park” (Souza et al., 2012) – resides, in the presence of taxa related to marine lineages (Rebollar et al., 2012; Souza et al., 2006; Souza, Eguiarte, et al., 2018), rich prokaryotic communities (López-Lozano et al., 2013; Pajares et al., 2012, 2013; Souza, Olmedo-Álvarez, et al., 2018), and diverse fungal (Velez et al., 2016) and viral communities (Taboada et al., 2018), as well as a high microbial diversity within the microbial mats and stromatolites (Bonilla-Rosso et al., 2012; de Anda et al., 2018; Peimbert et al., 2012).

In March 2016, we discovered an outermost elastic layer microbial mat in CCB, which enables the formation of dome-like structures under wet conditions (Fig. 1). The internal microbial layer, insulated by an outer aerobic community, thrives under an anaerobic environment, rich in methane, hydrogen sulfide, and small volatile hydrocarbons that recreate the atmosphere of the Archaean Eon, hence we named these mats “Archaean Domes” (Medina-Chávez et al., 2020). Similar types of elastic mats have previously been described in marine tidal zones, although their formation is relatively rare (Gerdes et al., 1993).

**Figure 1.**
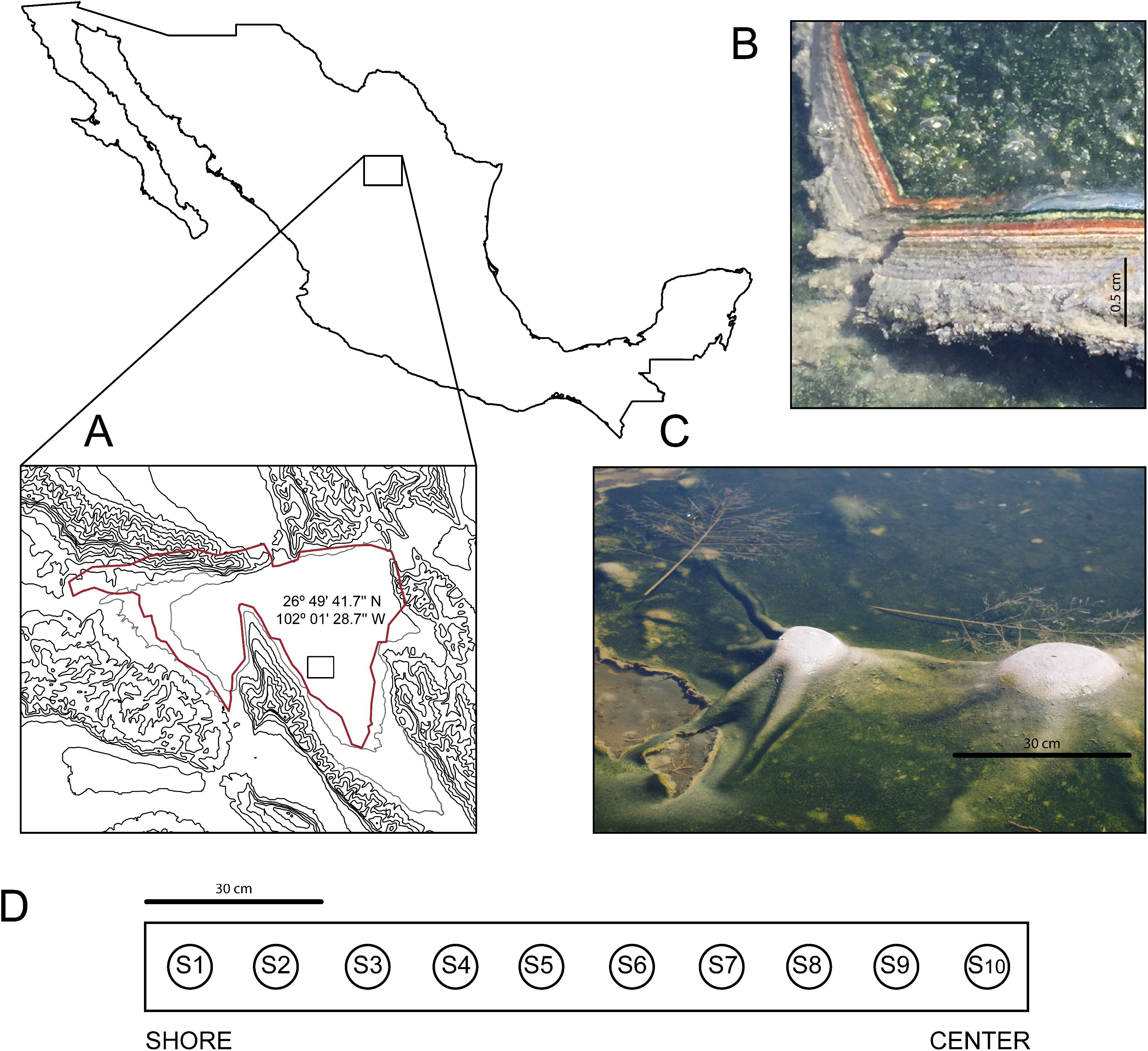
Site and sample (A) The Cuatro Ciénegas Basin within México and geographic location of the Archaean Domes in Pozas Azules Ranch. (B) Details of an Archaean Domes slice. (C) Domes when the sampling was conducted (note the size scale). (D) Sampling design along border toward center 1.5m transect (sample 1 (S1) to sample 10 (S10). Photo credit: David Jaramillo

Microbial mats are ideal systems for the study of microbial community structure under extreme conditions (Paerl & Yannarell, 2010; Prieto-Barajas et al., 2018). In this sense, there are approaches that can improve our understanding of microbial mat communities, such as determining the phylogenetic diversity of communities under extreme environmental conditions where, for example, abiotic filtering and competitive exclusion have been reported as drivers of phylogenetic clustering (Goberna et al., 2014;); another possible approach is to describe microbial subcommunities, e.g. those of rare taxa, a less-studied topic than the whole community (Liu et al., 2019) but nevertheless important to explore (Jia et al., 2018); other form is to analyze the metabolome of communities, which is one of the functional expressions of diversity and an underexplored aspect of microbial communities that can help complete our understanding of the observed diversity.

The investigation of microbial mats has gained special relevance in recent years with regards to the interpretation of structuring processes in microbial communities. However, hypersaline elastic mats remain poorly studied, limiting our understanding of microbial diversity and functioning under extreme environmental conditions (e.g. high salinity concentrations, elevated evaporation and frequent desiccation cycles). In this study, we evaluated microbial diversity and metabolome profile within this elastic hypersaline microbial mat from CCB, across ten samples collected on a 1.5-meter transect. Specifically, we aimed: 1) to explore microbial diversity by analyzing amplicons of the 16S rDNA and ITS regions, 2) to determine whether rare (low read count) and abundant (high read count) taxa are phylogenetically structured; and 3) to evaluate microbial metabolome.

## Materials & Methods

### Sampling site and sample collection

During the dry season (i.e. most of the year at CCB; Montiel-González et al., 2018), the microbial mats of the Archaean Domes remain humid under a salty crust, which is then dissolved during the rainy season, allowing for the development of the dome-like structures.

The microbial mats of the Archaean Domes were sampled during the rainy season of 2018 (October), with ten evenly spaced samples (S1-S10) along a 1.5-meter transect, from border toward center, (26° 49’ 41.7’’ N 102° 01’ 28.7’’ W) in the ‘Pozas Azules’ ranch of Pronatura Noroeste within CCB (Fig. 1). Microbial mats were collected and transferred to sterile conical tubes (50 mL), stored at 4°C, and subsequently frozen in liquid nitrogen until processing in the laboratory. Sampling was under the collection permit SGPA/DGVS/03188/20 issued by Subsecretaría de Gestión para la protección Ambiental, Dirección General de Vida Silvestre.

### Total DNA extraction and amplicons sequencing

Prior to DNA extraction, approximately 0.5 g of each mat sample was rinsed with sterile water, removing associated sediments. Total DNA extraction was performed following the protocol reported in de Anda et al. (2018).

DNA was sequenced at the Laboratorio de Servicios Genómicos, LANGEBIO (http://langebio.cinvestav.mx/labsergen/) for the V3 region of the 16S rDNA gene (357F universal primer 5’-CTCCTACGGGAGGCAGCAG-3’ / 519R universal primer 5’- GWATTACCGCGGCKGCTG-3’; (Lane 1991), and the ITS DNA region (ITS1F 5’- TCCGTAGGTGAACCTGCGG- 3’ / ITS4R 5’-TCCTCCGCTTATTGATATGC-3’ for ITS; (White et al., 1990). For each sample, 3 ug of genomic DNA (DO 260/280 1.8) was used; 16S amplicons were sequenced within Illumina Next-Seq 500 platform (2×150 PE, using forward reads; reverse reads were discarded from the analysis due to low quality), and the ITS amplicons were sequenced with a Mi-Seq platform (2×300 PE).

### Bioinformatics analyses

The 16S rDNA amplicon reads were quality filtered and dereplicated using the QIIME 2 bioinformatics platform version 2019.7 (Bolyen et al., 2018). Divisive Amplicon Denoising Algorithm 2 (dada2) denoise-single was used; sequences were demultiplexed and truncated at the 20^th^ bp from the left, and the 120^th^ bp from the right. ASVs were then taxonomically classified using the silva_132_99_v3v4_q2_2019-7 database. For all diversity analyses the final ASV table was rarefied to 760,000 reads per sample.

The ITS amplicon reads were processed using the ITS-specific workflow of the *dada2* v1.13.1 package (Callahan et al., 2016) in R (R Core Team, 2020). Briefly, *cutadapt* was used to remove primer and adapter contamination from the raw reads (Martin, 2011). Prior to determining ASVs, reads with inconsistent bases were excluded, and reads were required to have less than two expected errors based on their quality scores. ASVs classification was inferred from forward and reverse reads using the run-specific error rates.

Due to the wide range of sizes of the ITS amplicons reported for fungal species in CCB (485-782 bp; (Velez et al., 2016)), two different strategies were implemented to define taxonomic diversity. First, the post-denoised ASVs were merged after excluding reads with an overlap of less than 12bp and reads with mismatches in the overlapping region. Chimeras were removed using the consensus method of “removeBimeraDenovo” implemented in dada2. The taxonomic assignment was performed with the UNITE database (Nilsson et al., 2019), using the RDP naive Bayesian classifier (Wang et al., 2007) available in the *dada2* package with a minimum bootstrap value of 80. ASVs with a minimum of 85% identity and with E-values lower than e-45 were retained (Merge ITS data: Suppl. Table 1). ASVs without taxonomic assignment (mostly non-fungi organisms) were classified using *blastn* (non-redundant nucleotide collection (nr/nt) database) (Altschul et al., 1990), where only those ASVs within identities greater than 90% were enlisted (Suppl. Table 2). The remaining ASVs were considered non-classified.

As a great proportion of the reads failed to merge (∼56%), a parallel classification was implemented. The RDP naïve Bayesian classifier was used on the two sets of reads (forward and reverse) to check for congruence in the classification between the ASVs. ASVs that had an assignment bootstrap value lower than 50, or showed inconsistent classifications among forward and reverse reads, were aligned to the NCBI database (non-redundant nucleotide collection (nr/nt) database), to corroborate their taxonomic assignment. A more robust classification of ASVs was reached when using forward ASVs, thus all subsequent analyses were performed using the Forward-Only ASVs; even though some ASVs reached a genus level assignment, the taxonomic classification of forward reads was limited to the phylum and kingdom level (ITS table on Suppl. Table 3 and Suppl. Table 4); the Eukaryotic classification followed the UNITE database. The ITS community composition table (Forward-Only data) was rarefied to 164,820 reads using the *vegan* package (Oksanen et al., 2013) in R (R Core Team, 2020).

In sum, the filtered 16S rDNA amplicon sequences included 760,326–1,973,982 high quality reads per sample, with a total of 6,063 ASVs (Suppl. Table 5). The filtered ITS amplicon sequences included 164,822–339,394 high quality reads per sample, with a total of 923 ASVs (Suppl. Table 4). ITS amplicons were obtained for only six samples (S1, S2, S3, S4, S6 and S7) due to low DNA amount that remained in four samples (S5, S8, S9 and S10), after 16S rDNA amplicon sequencing.

### Alpha and beta diversity estimates

To evaluate all samples from 16S (S1-S10) and 6 samples (S1-S4, S6, S7) from the Forward-Only ITS data, we generated rarefied diversity plots using the *vegan* package (Oksanen et al., 2013) in R (R Core Team 2020).

In order to visualize and count the number of overlapping ASVs within the ten samples from the 16S rDNA gene and the six samples from the ITS region, we constructed an UpSet plot using the *UpSetR* package (Conway et al., 2017) in R (R Core Team 2020). Alpha diversity (Shannon index) and beta diversity (Bray-Curtis, Jaccard distances) were estimated using the 16S rDNA ASV community composition table (Suppl. Table 5), as implemented in Qiime2. For the Forward-Only ITS ASV community composition table (Suppl. Table 4), alpha and beta diversity were estimated with the *phyloseq* package (McMurdie & Holmes, 2013) in R (R Core Team, 2020). The beta diversity distance matrix was used to construct an UPGMA dendrogram of samples (Carteron et al., 2012). Mantel tests (999 permutations) (Legendre & Legendre, 2012) were conducted with the *vegan* package (Oksanen et al., 2013) to unveil the relationships between beta diversity and geographic distance, as well as between beta diversity and the metabolomic profile distance matrix (see below).

### Phylogenetic diversity

We estimated phylogenetic diversity within samples using Faith’s Phylogenetic Diversity index (Faith, 1992) and the Phylogenetic Species Clustering index (Helmus et al., 2007) using the *picante* package (Kembel et al., 2010) in R (R Core Team 2020). Given a phylogenetic tree, phylogenetic diversity (PD) estimates the total length of branches encompassing a subset of taxa (*e.g.*, sampled communities), where PD increases as the subset of taxa is more phylogenetically diverse (Faith, 1992). Phylogenetic Species Clustering (PSC) is a measure of the degree of clustering of taxa across the phylogeny (Helmus et al., 2007), where a PSC value approaching zero indicates that the taxa analyzed were phylogenetically clustered.

To estimate the two metrics, we used the rarified diversity matrices and the corresponding phylogenetic trees for bacterial and fungal ASVs, separately. We tested the observed patterns of PSC across samples against 1,000 replicates under a null model and estimated standardized effect sizes (SES) for this index (null model Phylogenetic Species Clustering, nmPSC). This allowed us to identify samples that had more or less phylogenetic clustering than expected by chance alone. To construct the null model, we randomized the entries of the rarified diversity matrices while keeping total sample diversity constant. In addition, we estimated PD and PSC for the most abundant and rare ASVs within each site. To define rare (low proportionated or low read count) and abundant (high proportionated or high read count) ASVs, we estimated the first and third quartiles (1Q, 3Q) of the distribution of ASV (organized by low to high read count), for each site separately. We used the sites’ 1Q and 3Q as the thresholds to define rare and abundant ASVs; the sites’ 1Qs ranged from 3 to 16 reads per ASV, whereas the 3Qs ranged from 37 to 306 reads per ASV; rare and abundant ASV numbers using this method were equivalent per site. We estimated SES for PD and PSC as defined above, using matrices for rare and abundant ASV. The phylogenetic tree was generated by QIIME 2 (Bolyen et al., 2018) for 16S data. For the ITS data (Forward-Only data), a phylogeny was obtained by applying an equal phylogenetic QIIME 2 methodology, using Maftt (Katoh & Toh, 2010) and FastTree (Price et al., 2009) within Cipres Science Gateway (Miller et al., 2010) with default XSEDE parameters.

### Metabolomics

#### Soluble compounds extraction and LC-MS/MS analysis

The samples (10 g each) were extracted with 60 mL of CHCl_3_-MeOH (1:1) in an orbital shaker at 150 rpm for 24 hrs. The mixture was filtered, and the solvent was evaporated under reduced pressure. The dried extracts were reconstituted in 60 mL of CH_3_CN-MeOH (1:1) and defatted with hexanes. Then the CH_3_CN-MeOH layer was dried under a vacuum and resuspended in MeOH (LC-MS grade) to yield a concentration of 1 mg mL^−1^. All samples were filtered with a 0.22 μm membrane and analyzed using a Waters Acquity Ultraperformance Liquid Chromatography (UPLC) system (Waters Corp.) coupled to a Thermo Q Exactive Plus (ThermoFisher) mass spectrometer. The UPLC separations were performed using an Acquity BEH C_18_ column (50 mm × 2.1 mm I.D., 1.7 μm; Waters) equilibrated at 40 °C and a flow rate set at 0.3 mL min^−1^. The mobile phase consisted of a linear CH_3_CN-H_2_O (acidified with 0.1% formic acid) gradient starting at 15% to 100% of CH_3_CN over 8 min. The mobile phase was held for another 1.5 min at 100% CH_3_CN before returning to the starting conditions. High-resolution mass spectrometry (HRMS) data and MS/MS spectra were collected in the positive/negative switching electrospray ionization (ESI) mode at a full scan range of *m/z* 150−2000, with the following settings: capillary voltage, 5 V; capillary temperature, 300°C; tube lens offset, 35 V; spray voltage 3.80 kV; sheath gas flow and auxiliary gas flow, 35 and 20 arbitrary units, respectively.

### Untargeted metabolomics

The HRMS-MS/MS raw data was individually aligned and filtered with MZmine 2.17 software (http://mzmine.sourceforge.net/) (Pluskal et al., 2010). Peak detection was achieved as follows: *m/z* values were detected within each spectrum above a baseline, a chromatogram was constructed for each of the *m/z* values that spanned longer than 0.1 min, and finally, deconvolution algorithms were applied to each chromatogram to recognize the individual chromatographic peaks. The parameters were set as follows for peak detection: noise level (absolute value) at 1×10^6^, minimum peak duration 0.5 s, tolerance for *m/z* variation 0.05, and tolerance for *m/z* intensity variation 20%. Deisotoping, peak list filtering, and retention time alignment algorithm packages were employed to refine peak detection. The join align algorithm compiled a peak table according to the following parameters: the balance between *m/z* and retention time was set at 10.0 each, *m/z* tolerance at 0.05, and retention time tolerance size was defined as 2 min. The spectral data matrix (comprised of *m/z*, retention time, and peak area for each peak) was imported to Excel (Microsoft) for heatmap analysis using *R* (R Core Team, 2020).

Molecular networking was performed from the mzML files using the standard Global Natural Products Social (GNPS) molecular networking platform workflow with the spectral clustering algorithm (http://gnps.ucsd.edu; Aron et al, 2020). For spectral networks, parent mass of 0.01 Da and fragment ion tolerance of 0.02 Da were considered. For edges construction, a cosine score over 0.70 was fitted with a minimum of 4 matched peaks, and two nodes at least in the top 10 cosine scores (K) and maximum of 100 connected the components. After that, the network spectra were searched against GNPS spectral libraries, and graphic visualization of molecular networking was performed in Cytoscape 3.8.0 (Shannon et al., 2003). Chemical structural information within the molecular network was obtained using the GNPS MolNetEnhancer workflow (Ernst et al., 2019), which incorporated *in silico* structure annotations from the GNPS library search (metabolomic_support_info.zip in supplemental files). The chemical profiles were also manually dereplicated using UV-absorption maxima and HRMS-MS/MS data against the Dictionary of Natural Products v 29.1 and Dictionary of Marine Natural Products 2019 (Taylor and Francis Group), MarinLite (University of Canterbury, New Zealand) and SciFinder (CAS) databases as described by El-Elimat et al. (El-Elimat et al., 2013), for fungal and bacterial small molecules. For candidate search, exact mass accuracy was set to 5 ppm (Sumner et al., 2007). Heatmap visualization of metabolomic data table was generated in MetaboAnalyst (Pang et al., 2020).

## Results

### Microbial taxonomic composition

Overall, 40 prokaryote phyla were detected from the 16S rDNA gene libraries (Suppl. Fig. 1A), and a total of 6,063 different ASVs were found, with a total proportion of 11.8% of unclassified reads (Suppl. Table 3). Bacteroidetes was the most abundant (high read count) phylum, representing around 23% of all sequences in the samples, followed by Proteobacteria with ≈ 19% and Cyanobacteria with nearly 18%. Spirochaetes and Chloroflexi represented ≈ 8% and ≈ 4%, respectively. Nearly 2% were Firmicutes, Patesibacteria and Halanaerobiaeota (Suppl. Fig. 1A).

Genera composition is shown in Fig. 2. A total of 219 genera of Bacteria were detected in the 10 sampled sites (S1-S10) at a 1.5 m-scale. From those, only 9 genera had high read counts (representing ≈ 80% of the total proportion, Fig. 2 A bottom): *Coleofasciculus* (formerly *Microcoleus chtonoplastes*), *Spirochaeta*, *Catalinimonas*, *Desulfovermiculus*, *Halanaerobium*, *Tangfeifania*, *Sungkyunkwania*, *Sediminispirochaeta* and *Imperialibacter*. The remaining 210 genera can be classified as rare biosphere, with ≈ 20% of the total proportion (less than 1% of the total proportion each, Fig. 2 A, top), having a heterogeneous distribution between sites. As for 16S rDNA ASVs, 18 of the 6,063 were abundant (>1% relative abundance each), representing 49% of the relative abundance of reads (Suppl. Table 5). This 18 ASVs were distributed within the previously mentioned highly abundant (high read count) genera and phyla (except Firmicutes and Patesibacteria).

**Figure 2.**
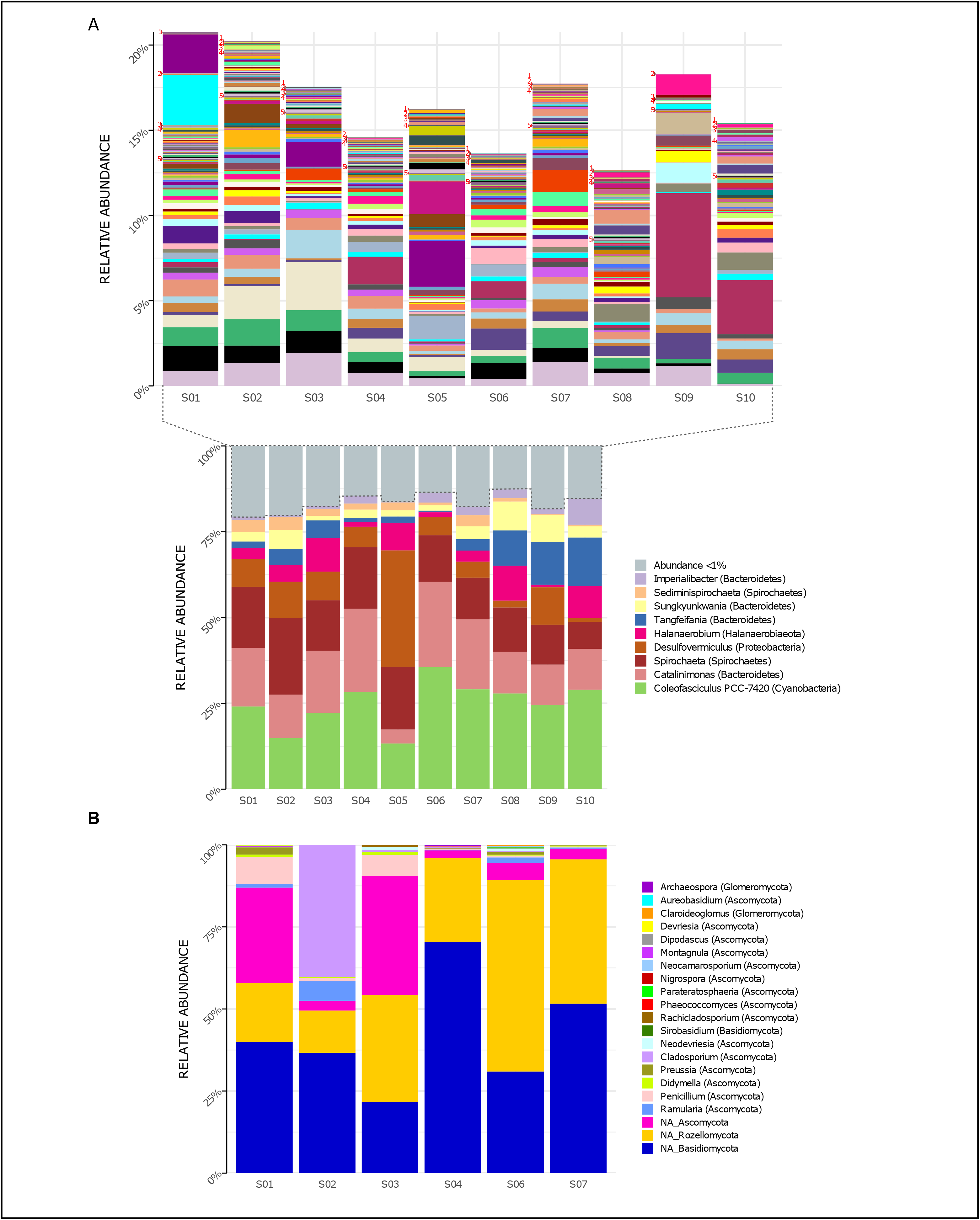
Genera relative proportion of taxa in Archaean Domes, Cuatro Ciénegas Basin sample A) Bacteria (bottom: shows the most common taxa, more than 1% of total proportion of taxa; top: shows less than 1% of total proportion; for labels of 210 corresponding genera, check Supplementary Table 10) B) Fungi (using Merge data, only fungal ASVs. NA is for ASVs that were not assigned to any genera, but phylum only)

Conversely, 6,045 ASVs were rare taxa (<1% relative abundance each) which represents 51% of total reads: 4,124 of them were non-classified, and the rest (1,921) were distributed within the whole 40 bacterial phyla and almost all (217) genera (Suppl. Table 5). The two exceptions of genera with no rare ASVs were *Tangfeifania* with 2 highly proportionated ASVs, and *Coleofasciculus* with only one high-read count ASV. Moreover, 28 ASVs of Archaea were obtained using the 16S primers, comprising two phyla, Woesearchaeota and Euryarchaeota (Suppl. Fig. 1A). We detected 4 genera within Halobacteria class: *Halovivax*, *Halorubrum, Halorussus* and *Haloasta*.

For the ITS region libraries, we successfully obtained reads for six of the sampled sites (S1 to S4, S6 and S7). For the Forward-Only ITS data, we detected 923 different ASVs (Suppl. Table 4). Many of the classified reads belong to Alveolata protists (77%, Suppl. Fig. 1B), followed by Fungi (11%) and Viridiplantae (8%); Metazoa, Protista, Amoebozoa, Rhodoplantae, Chromista and Stramenophila comprised the remaining 4%. Unassigned taxa were 50% of all reads (Suppl. Table 3). The majority of ASVs within Forward ITS data could not be accurately classified at lower taxonomic categories, particularly for three common ASVs (ASV0 (unclassified), ASV1 (Ciliophora) and ASV2 (Ciliophora)), which comprise nearly 70% of the proportion of ITS reads for all sites.

Using Merge ITS ASVs data (Suppl. Table 1) for a better taxonomic assignment, Fungal ASVs comprising 18 genera were detected, with phyla Ascomycota, Basidiomycota and Rozellomycota present in all samples (Fig. 2B). Non-fungal Eukarya Merge ITS sequences classification (hits <90%) are shown in Suppl. Table 2*. Fabrea salina* (Alveolata), and *Vannella simplex* (Amebozoa) were two species that constituted nearly 0.5% of the total proportion of Merge ITS ASVs data, among other classified but low-read count eukaryotic ASVs.

### Alpha and beta diversity

Rarefaction curves (Suppl. Fig. 2A and 2B) showed similar results for both the Forward-Only ITS region libraries (S1, S2, S3, S4, S6, S7), and the 16S rDNA gene libraries, as all sites reached an asymptote, suggesting an adequate sampling effort. Only 183 ASVs (3% of the total ASV read count) were shared between all sites for 16S rDNA gene libraries (Fig. 3A), forming a very small core, but comprised 72.4% of all reads (composed of twenty-one phyla), while most of the ASVs were unique to a specific site or shared between two or three sites. For ITS libraries, the 56 shared ASVs (6%) comprised 94.7% of all reads (Fig. 3B).

**Figure 3.**
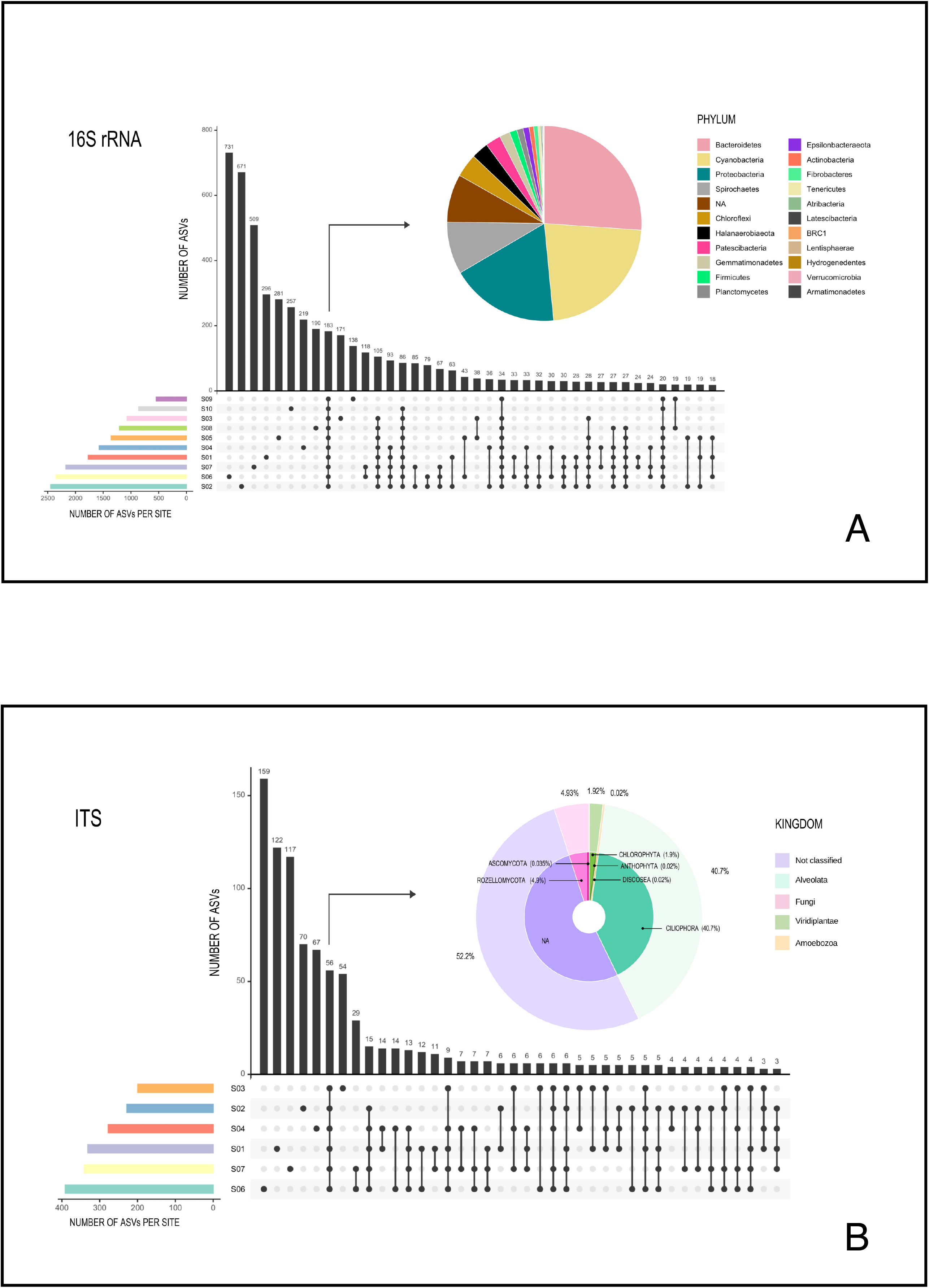
Graphic representation of shared ASVs within Archaean Domes sampling sites Number of shared ASVs is represented by black bars, showing the exact number on the top. Dots below bars denote sites where those ASVs are shared, and lines joining dots indicate the shared sites. Colored bars at bottom left of the main figure are for ASVs number within each site. Pie charts show taxonomy of the few shared ASVs (corresponding bars are pointed with arrows) within all sites. All of these shared ASVs had a high proportion within the whole ASVs (72.5% of 16S rDNA data and 95% of Forward-Only ITS data). (A) 16S Archaea and Bacteria, rDNA gene data; (B) Fungi and other Eukarya, Forward-Only ITS data

The 16S rDNA ASVs alpha diversity (Table 1) ranged from 2,399 observed ASVs in S2 (Shannon = 7.4, highest Shannon index value), to 544 observed ASVs in S9 (Shannon = 6.1). In the case of the ITS region libraries (Forward-Only ITS data), the highest ASV number for Eukarya was S6 with 383 ASVs and a Shannon diversity of 2.5; S3 had the lowest observed ASVs with 199, but this site had the highest Shannon value of 3 (Table 1).

**Table 1.** Alpha diversity indices Alpha diversity indices for 16S rDNA data and Forward-Only ITS region data from Archaean Domes CCB microbial mats

| ITS data |  |  |  |  |  |
| --- | --- | --- | --- | --- | --- |
| $\alpha$ diversity | | | Phylogenetic diversity | | |
| Sample | Observed | Shannon | Faith PD | Faith PD rare | Faith PD abundant |
| 1 | 322 | 1.827 | 86.309 | 42.596 | 18.161 |
| 2 | 226 | 2.713 | 60.622 | 40.846 | 14.516 |
| 3 | 199 | 3.062 | 47.289 | 30.370 | 14.774 |
| 4 | 270 | 1.783 | 83.008 | 51.513 | 16.287 |
| 6 | 383 | 2.501 | 131.435 | 75.352 | 28.794 |
| 7 | 337 | 2.008 | 114.983 | 60.352 | 18.865 |

| 16S rDNA data |  |  |  |  |  |
| --- | --- | --- | --- | --- | --- |
| $\alpha$ diversity | | | Phylogenetic diversity | | |
| Sample | Observed | Shannon | Faith PD | Faith PD rare | Faith PD abundant |
| 1 | 1761 | 6.863 | 544.704 | 290.491 | 44.159 |
| 2 | 2399 | 7.402 | 789.807 | 368.430 | 84.442 |
| 3 | 1068 | 6.256 | 211.745 | 156.204 | 20.064 |
| 4 | 1564 | 6.499 | 500.787 | 259.828 | 40.627 |
| 5 | 1344 | 6.882 | 220.536 | 172.299 | 21.557 |
| 6 | 2345 | 6.301 | 851.168 | 302.103 | 131.337 |
| 7 | 2156 | 6.838 | 759.536 | 328.238 | 97.486 |
| 8 | 1195 | 6.487 | 205.987 | 139.839 | 22.798 |
| 9 | 544 | 6.126 | 160.368 | 94.201 | 18.577 |
| 10 | 828 | 5.115 | 283.017 | 148.655 | 36.514 |

Pairwise dissimilarities between samples for both the 16S rDNA gene and ITS region data ranged from 0.2 to 0.7 (16S) and 0.2 to 0.6 (ITS) for the Bray-Curtis distance values and 0.5 to 0.8 (16S) and 0.3 to 0.7 (ITS) according to the Jaccard-1 values (Suppl. Fig. 3 A and B). For both distance values, S10 is different from the rest of the 16S rDNA gene libraries, and S1, S2, S4, S6, S7 were associated in the dendogram clusters. In the ITS libraries, there are two main clusters (S1, S2 and S4; S3, S6 and S7). A significant geographic pattern in the Mantel test (*p*=0.0402, r=0.3224) was found for Jaccard-1 of the 16S library, however we did not find significant values based on geography for ITS libraries for either Jaccard-1 or Bray-Curtis (Suppl. Table 6).

### Phylogenetic diversity

For both libraries, Faith Phylogenetic Diversity indicated that rare (low read count) ASVs on each site, were more diverse than abundant (high read count) ASVs (Table 1). In respect to the null model distribution of Phylogenetic Species Clustering (nmdPSC), the 16S and ITS communities analyzed per site had in some sites a phylogenetic clustering distribution, and in others sites a null model distribution compared with chance alone (Fig. 4 A and B, dots between lines or below -2).

**Figure 4.**
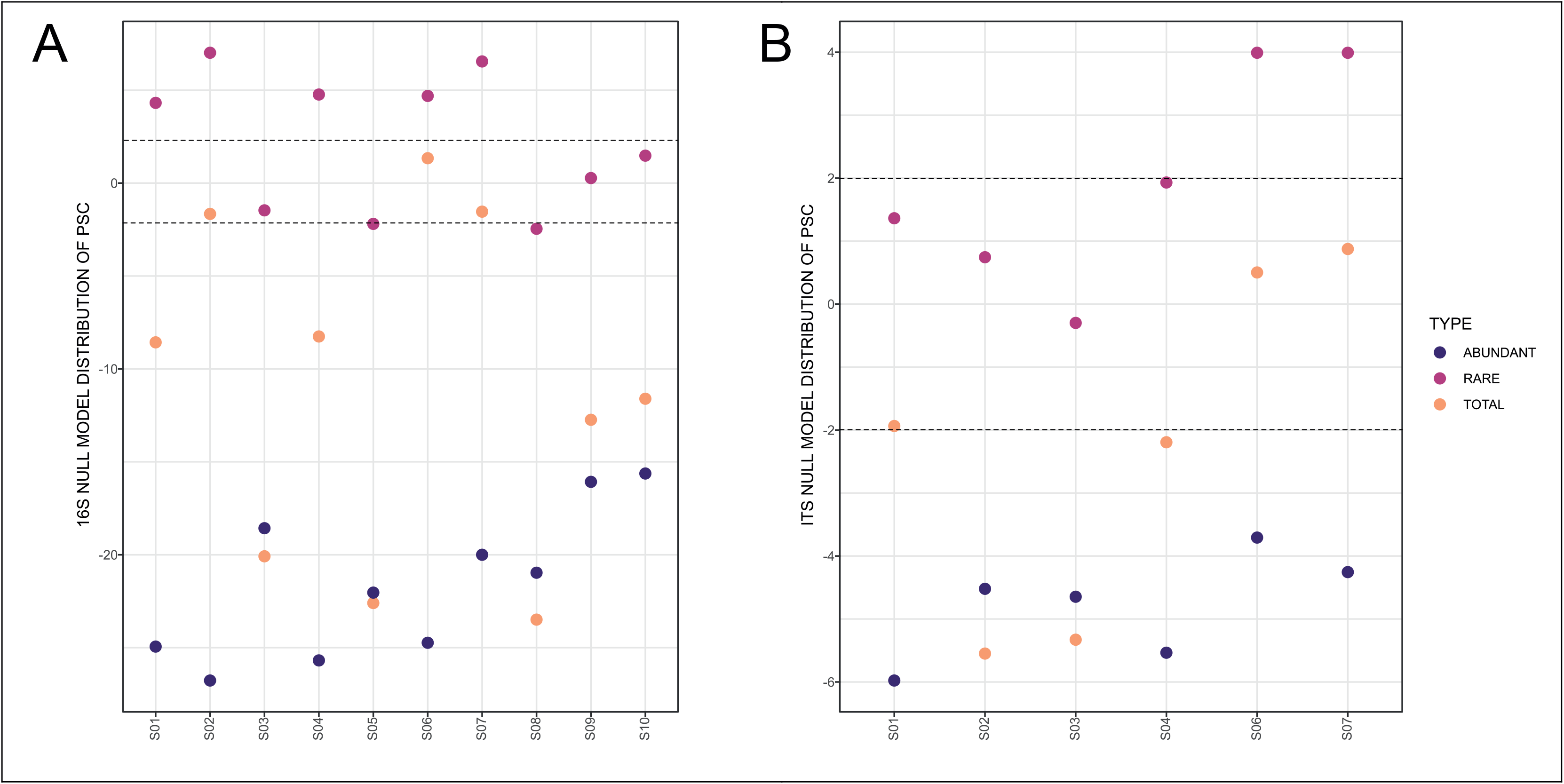
Graph representing null model distribution of Phylogenetic Species Clustering (nmdPSC) (A) 16S Archaea and Bacteria, rDNA gene data; (B) Forward-Only ITS data. Dotted lines are showing nmdPSC significant values: > +2 is consistent with a sample that had more phylogenetic overdispersion than expected by chance alone, and nmdPSC value < -2 indicates more phylogenetic clustering than expected by chance alone; values in the range +2 and -2 had a null model distribution. nmdPSC values are reported in Suppl. Table 7.

When analyzing rare and abundant subcommunities using an equivalent ASV number per site, nmdPSC indicated that abundant ASVs were phylogenetically clustered within all sites and for both libraries, while rare ASVs were subcommunities with a null model distribution in some sites, or phylogenetically dispersed in other sites, for bacterial and fungal libraries (Fig. 4 A and B, Suppl. Table 7).

### Metabolomic profile

Based on the LC-MS results of S1-S10, a wide range of metabolites were visualized on a heatmap analysis chart (Suppl. Fig. 4 A&B) and a GNPS molecular networking analysis (Fig. 5 and Suppl Table 4).

**Figure 5.**
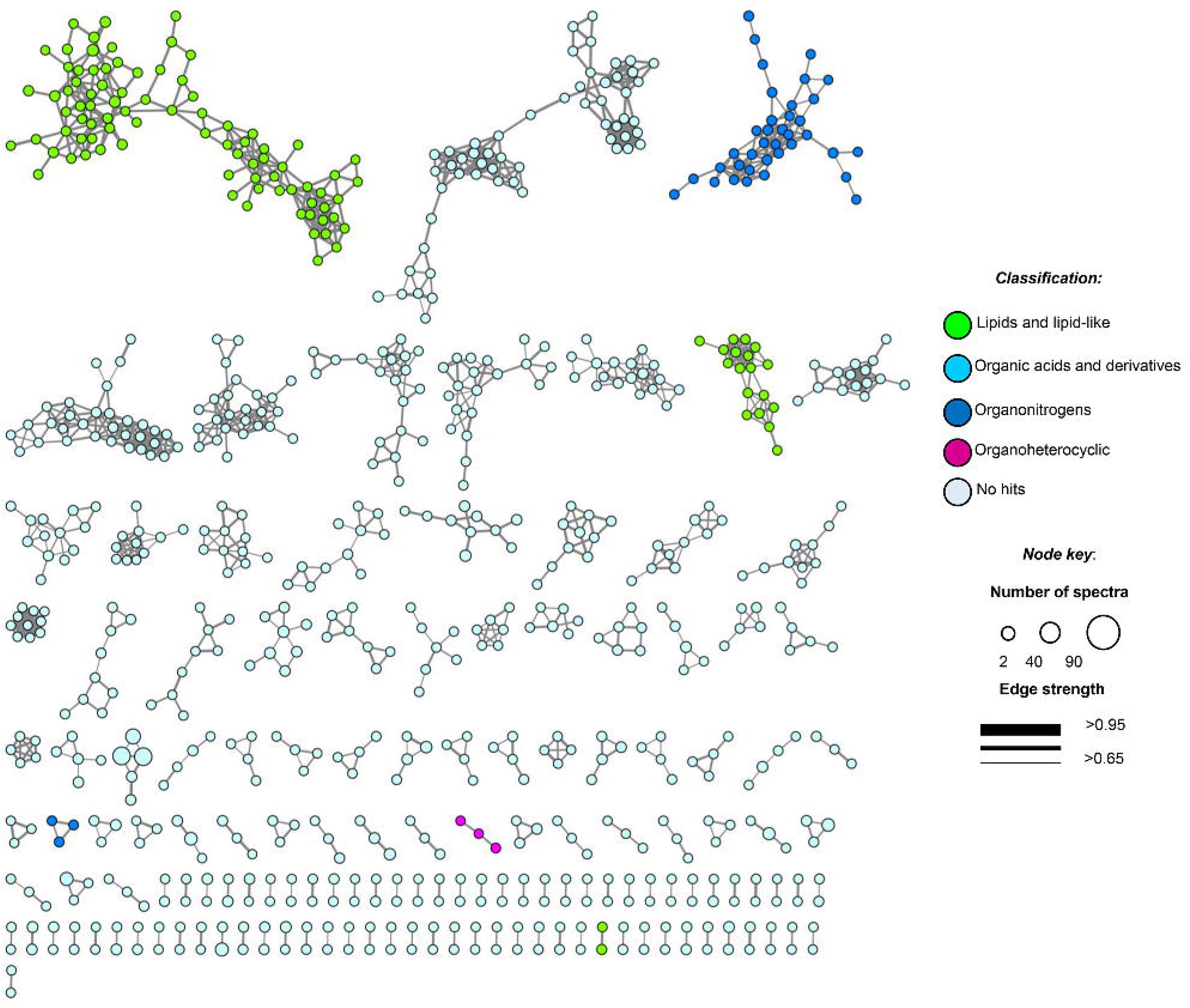
Molecular network of soil samples grouped metabolite features into 139 chemical families Each node in the network represents one metabolite feature. Nodes connected to each other are structurally-related (cosine MS2 similarity score ≥0.7)

Untargeted metabolome analysis of microbial communities in soil or environmental samples is difficult, due to the complexity and heterogeneity of this matrix and the scarcity of the secondary metabolites extracted. In the heatmap, the 1,341 molecular features at unique retention times (Suppl. Fig. 4 A&B) shared several common regions across all samples (molecular features), allowing us to group them into three main clusters (cluster 1, samples 3-5 and 10; cluster 2, samples 1, 2 and 6; and cluster 3, samples 7-9, Suppl. Fig. 4B). We found highly heterogeneous metabolic profiles in all samples, and no significant associations were observed between metabolite features and ITS or 16SrDNA community composition matrices (Suppl. Table 6).

Moreover, feature-based GNPS molecular networking analysis grouped the metabolite features in 1,534 nodes distributed into 67 chemical families (sub-networks) of ≥3 members, 72 families of two metabolite features, and 765 singletons (Fig. 5). Chemical ontology analyses revealed the presence of superclasses of lipids and lipid-like compounds, organic acids, and organonitrogen and organoheterocyclic compounds, all commonly found in soil samples (Fig. 5). Detected compounds in the GNPS and manual metabolome analysis of the main ions observed in LC-MS/MS data focused on microbial metabolites, allowing tentative identification of several secondary metabolites that passed our quality criteria (≤ 5 ppm mass error; Suppl. Table 8). Finally, some plant-derived secondary metabolites and small molecules reflecting human activity were also observed (Suppl. Table 8).

## Discussion

### Microbial composition at a small scale

Our study describes microbial diversity on a 1.5 m scale in a rare elastic microbial mat located at an extreme site. Altogether, we found 40 prokaryotic phyla, including 219 genera of Bacteria and 4 of Archaea, as well as 18 Fungal genera (Fig. 2 and Suppl. Fig. 1).

#### Bacteria

The bacterial phyla (Suppl. Fig. 1A) detected in our study site are common components of hypersaline microbial mats (Bolhuis et al., 2014; Visscher et al., 2010): Bacteroidetes, Proteobacteria, Cyanobacteria, Chloroflexi, and Firmicutes, among other abundant (high read count) phyla, which are similar at this broad taxonomic level to other sites in CCB (Bonilla-Rosso et al., 2012; Breitbart et al., 2009; de Anda et al., 2018; Nitti et al., 2012). As expected from a microbial mat, a primary producer was present and abundant in all of our samples: *Coleofasciculus* (Fig. 2), a filamentous nitrogen-fixing cyanobacteria genus well-adapted to saline conditions (De Wit et al., 2013; Ramos et al., 2017) that has been reported in Guerrero Negro microbial mats (Des Marais, 2010; Wong et al., 2016). We also found the abundant taxon *Catalinimonas,* which is a halophilic Bacteroidetes marine genus (Choi et al., 2013) that has not been previously reported in microbial mats. Another common taxon in our samples is the *Spirochaeta* genus (Bolhuis et al., 2014; Margulis et al., 2006), an anaerobic chemo-organotrophic taxon abundant in hypersaline marine mats (Ben Hania et al., 2015). Noteworthy is that the *Spirochaetes* phylum has not been detected previously in any CCB mats (Bonilla-Rosso et al., 2012; de Anda et al., 2018; Peimbert et al., 2012). We also observed *Desulfovermiculus*, a sulfate-reducing Deltaproteobacteria, and *Halanaerobium*, a thiosulfate-reducing Firmicute that is halophilic and can form spores (Liang et al., 2016), both genera are common in marine microbial mats (Bolhuis et al., 2014), reinforcing previous evidence from CCB studies that this site can harbor taxa that are phylogenetically similar to marine lineages (see Rebollar et al., 2012; Souza et al., 2006, 2018).

#### Archaea

Archaea data presented here should be taken cautiously, since the primers for 16S rDNA used are universal but were not designed specifically to amplify archaea and would likely recover only 1% or less of archaeal diversity. However, we decided to only describe the groups detected, avoiding any interpretations regarding quantity where there could be a strong bias. We detected two main phyla (Suppl. Fig. 1A); these were Woesearchaeota, a phylum that potentially has a syntrophic interaction with methanogens (Liu et al., 2018; Ortiz-Alvarez & Casamayor, 2016), and Euryarchaeota, within which we found four halophilic genera: *Halovivax*, *Halorubrum, Halorussus* and *Haloasta*, all of them previously reported in saline habitats and sediments (McGonigle et al., 2019; Zaitseva et al., 2018).

In a recent metagenomic study of the Archaean Domes (Medina-Chávez et al., 2020), some archaeal taxa were detected in all sampled sites, forming a “core taxa”. In those metagenomes, Archaea represented nearly 5% of the total reads. The metagenomic data (Medina-Chávez et al., 2020) and the 16S rDNA gene amplicons were sampled from the same pond at different times and locations, and both contained these lineages common to hypersaline habitats.

#### Fungi

Fungal members represent a diverse and common component of hypersaline water columns and mats (Cantrell et al., 2013) and have been suggested to play a role in the degradation of EPS (exopolymeric substances) (Cantrell & Duval-Pérez, 2013). Our results suggest that unidentified uncultured members of the Ascomycota and Rozellomycota comprise the core mycobiota of the analyzed mats, as they occurred across all the samples (Fig. 3B). In addition, it is feasible that these abundant heterotrophs occupy a niche at the micro-scale of microbial mats, where they can degrade the abundant EPS produced in the CCB microbial mats (Breitbart et al., 2009; de Anda et al., 2018; Peimbert et al., 2012). In particular, halotolerant taxa such as members of the genera *Aspergillus*, *Penicillium*, and *Nigrospora* have been repeatedly reported in microbial mats at other locations (Cantrell et al., 2013; Cantrell & Duval-Pérez, 2013), evidencing their adaptation to extreme conditions. Thus, it appears that, similar to other microbial mats, fungal members could occupy an EPS-degrader role at the micro-scale of the Archaean Domes (Cantrell et al., 2011).

#### Other eukaryotes

In our sample, there are three abundant (high read count) ASVs of Alveolata protists that represent 70% of the ITS sequences (Suppl. Fig. 1B, Forward-Only ITS data) in all sequenced sites (S1-S4, S6, S7). Although the ITS primer pair used in this study were originally designed for fungal amplification, it has been reported that there are potential matches with non-fungal eukaryotic sequences as well (Toju et al. 2012). Nevertheless, these results should be interpreted in a descriptive sense, considering that several eukaryotic taxa are not being detected due to primer bias.

Two of the three ASVs are Ciliophora (i.e. ciliates), which is the most diverse phylum of protists, with 3,500 described species, most of which are cosmopolites (Gao et al., 2016). It will therefore be interesting to analyze more molecular markers in the future to distinguish the protist lineages, as well as to microscopically study the protists to reach a closer taxonomic identification. Other Protozoa ASVs were observed at low numbers and belong to the Amoebozoa phylum, and the ancestral algae from the kingdom Chromista were also observed at a low proportion (Suppl. Fig. 1B).

Although research on halophilic protozoan is still young, it has made great progress in recent years (Harding & Simpson, 2018). Protozoa in general are grazing organisms, as is the case with *Fabrea salina*, one of the few classified low proportion ITS ASVs in our study (Merge data, Suppl. Table 2). This ciliate is present in microbial mats and feeds on *Cyanotece sp.*, a Cyanobacteria associated with algae mats (Carrasco & Perissinotto, 2012). Indeed, in hypersaline environments, most ciliates are known to be important components of the pelagic microbial food loop (Liu et al., 2016). Amoebae have also been observed as grazers (Hauer et al., 2001; Hauer & Rogerson, 2005; F. Post et al., 1983) on cyanobacteria in surface mud samples from a salt pond in Eilat, within the Salton sea in Southern California (Hauer & Rogerson, 2005). Even though the diversity of protists is low, their prevalence, along with the large amount of viruses found in metagenomic samples (28% of the metagenomes from Medina-Chávez et al., 2020), strongly suggests that predator-prey interactions (e.g. protist-bacteria, Saleem et al., 2013) could be very important for the dynamics of the system (Carreira et al., 2020).

### Alpha and beta diversity

The Shannon Index accounts for both species evenness and abundance; hence, it might not reflect the total richness if most of the found taxa are part of the rare biosphere. For example, Site 3 for the Forward-Only ITS data had the highest Shannon value, despite having the lowest ASV number (Table 1), due to its higher evenness distribution of taxa. Meanwhile, the rarefaction and observed diversity reflects the richness of the site (Suppl. Fig. 2A and 2B; Table 1).

Beta diversity analysis showed values that represent a large ASV turnover between sites despite the small scale (Suppl. Fig. 3), illustrated by the heterogenous composition observed within bacterial genera (Figure 2A top). Accordingly, the different domains analyzed (i.e., the bacterial 16S rDNA data or the Eukarya Forward only ITS data) had a distinctive association pattern. This turnover pattern should be taken cautiously, as ASVs turnover might a methodological artifact: beta diversity analysis uses data on relative abundances, based on compositional data; this means that the presence of abundant ASVs might “push” the rare ASVs below the detection threshold. Therefore, this pattern should be corroborated in a future time-series using metagenomic data, and additionally, a transcriptome approach would explore if the rare ASVs are an active part of this community. A statistically marginal but significant geographic association was found only for the bacterial 16S rDNA data (p=0.0402, r=0.3224); this suggests a geographic pattern that also should be explored with a more extensive sampling.

### Bacterial community comparison with other CCB layered microbial communities

We compared our data with four previously published metagenomes of layered microbial communities from the CCB: one microbialite from a large spring-fed river Rio Mesquites (Nitti et al., 2012), one stromatolite from a blue pool in Pozas Azules (Breitbart et al., 2009) and two microbial mats, one from a small and shallow pond called Lagunita in Churince (de Anda et al., 2018), and the other from a green pool in Pozas Azules (Bonilla-Rosso et al., 2012).

At the phylum level, we found seven taxa present in all these five microbial communities: Actinobacteria, Bacteroidetes, Chloroflexi, Cyanobacteria, Firmicutes, Planctomycetes and Proteobacteria. All of them shared among all sites on Archaean Domes mats and had a high read count (Fig. 3A).

Additionally, Bonilla-Rosso et al., (2012) reported 28 orders that were shared within the microbial mat from a green pool (Bonilla-Rosso et al. 2012), the stromatolite (Breitbart et al., 2009) and a coastal hypersaline non-thermophilic microbial mat from Guerrero Negro (Kunin et al., 2008). We found 18 of those 28 orders in the Archaean Domes community (Suppl. Table 9).

We also explored lower taxonomic levels but did not find a consistent pattern. The common pattern at higher taxonomic ranks reflects, as Bonilla-Rosso et al., (2012) mentioned, that layered microbial communities assemble in biogeochemical gradients related to defined ecological niches (e.g. photosynthesis, sulfate reduction, heterotrophy) according to their functional traits, rather than their species (Burke et al., 2011, Escalas et al., 2019).

### Phylogenetic diversity

The null model distribution of Phylogenetic Species Clustering (nmdPSC, Fig. 4) of whole bacterial and eukaryotic communities showed negative or null values, indicating a clustered pattern compared with the null model. Phylogenetic clustering is predicted to be evident in environments with poor nutrient availability (Mondav et al., 2017), such as the microbial mat conditions. Since little is known about patterns and processes of rare and abundant bacterial and eukaryotic taxa in microbial mats, we also analyzed these two microbial subcommunities.

#### Abundant (high read count) taxa

The nmdPSC pattern of clustering for the abundant ASVs (Fig. 4) could be similar to some soil bacterial communities that typically show a coexistence of phylogenetically close relatives (Bryant et al., 2009; Horner-Devine & Bohannan, 2006), which may be the product of abiotic filtering but also of competitive exclusion (Goberna et al., 2014). Competitive exclusion seems to be a likely explanation for the phylogenetic clustering observed for abundant bacterial and eukaryotic ASVs, as proposed by Goberna et al. (2014), since the presence of compounds with antimicrobial properties (check Metabolomic profile sub-section below) suggests high levels of competition among abundant microbes.

#### Rare (low read count) taxa

The rare biosphere, when analyzing equivalent numbers of ASVs within abundant taxa, was more phylogenetically diverse (Faith Pylogenetic Diversity index, Table 1), as well as more phylogenetically disperse (nmdPSC, Fig. 4) than the abundant taxa within these samples for both the 16S rDNA and ITS libraries, suggesting that environmental filtering is not occurring for the rare biosphere in this site.

The two analyzed subcommunities (abundant and rare) showed different phylogenetic dispersion patterns. The clustering pattern for abundant taxa --and also for the whole community--, suggests that the associated assembly processes are abiotic filtering or competitive exclusion (Wang et al., 2013; Mondav et al., 2017). In contrast, communities that show phylogenetic dispersion, such as the rare taxa of the Archaean Domes, could be more affected by stochastic processes such as dispersal and drift (Wang et al., 2013; Mondav et al., 2017). Thus, both processes, deterministic and stochastic could be occurring in the Archean Domes community.

These rare and abundant patterns described suggest that rare taxa could be a large reservoir of genetic diversity, having diverse metabolic functions within this microbial mat (Jia et al., 2018) and perhaps presenting collective adaptations to the extreme conditions. This is in correspondence with the unique metabolome profile showed for each site, despite the homogeneous abundant taxa. These ideas should be taken cautiously, since rare taxa may not be an active part of the community, as stated previously.

### Metabolomic profile

The dynamic and complex metabolomic profiles (Suppl. Fig. 4A&B, Fig. 5 and Suppl. Table 8) of the studied microbial mat was consistent with the complex microbial community observed in the metagenomic analysis (Medina-Chávez et al., 2020). Even though the expression of regulatory genes responsible for secondary metabolites production depends on several factors (e.g. nitrogen or phosphate starvation, stressors like heat, pH and damage to the cell wall, among others) (Baral et al., 2018), along with the lack of soil metabolomics studies and the solubility of the compounds for LCMS analysis, we were still able to tentatively annotate several classes of compounds with interesting biological properties (Fig. 5 and Suppl. Table 8). Many of these compounds were found to overlap between most of the samples: the antibiotics 8,9-dihydroindanomycin (Li et al., 2009), neoindanomycin (Rommel et al., 2011) described from *Streptomyces* species; the cytotoxic daryamide C (Asolkar et al., 2006), gibbestatin B (Kinashi & Sakaguchi, 1984) and furaquinocin H (Ishibashi et al., 1991), from *Streptomyces* strains; the germination inhibitor colletofragarone A1, from fungi *Colletotrichum fragariae* and *C. capsiciae* (Inoue et al., 1996); periconiasin G (Zaghouani et al., 2016) from *S. nitrosporeus* and *Periconia* sp. F-31 with free radical scavenger and weak anti-HIV activities; and phomamide and tomaymycin I form *Phoma lingam* (Ferezou et al., 1980) and *Nocardia* sp. (Tozuka et al., 1983) with no biological activity described. Other metabolites of interest were found in specific samples, for example the antifungal agents griseofulvin and dechlorogriseofulvin, produced by microorganisms of the genera *Penicillium*, *Aspergillus*, *Xylaria*, *Coriolus*, *Palicourea*, *Memnoniella*, *Khauskia*, *Carpenteles*, *Arthrinium*, *Nigrospora*, *Stachybotrys*, and *Streptomyces* (Atta-ur-Rahmanro, 2005; Kimura et al., 1992; Knowles et al., 2019; Park et al., 2006) were identified only in S7; the antimitotic agent nostodione A from the blue-green alga *Nostoc commune* (Kobayashi et al., 1994) commune in samples S2 and S6-S8; and the mycotoxins citreohybridone B (Kosemura, 2003), setosusin (Fujimoto et al., 1996), roridin E (Böhner et al., 1965) and ganoboninone A (Ma et al., 2015) in samples S8-S10. Interestingly, several observed ions (Fig. 5) do not match with any data reported and could represent potentially new secondary metabolites. Finally, plant- and animal-derived metabolites and compounds resulting from human activity (contaminants), were also annotated (Suppl. Table 8).

The association of compounds between fungi and osmotrophs, as well as to other Eukarya, reflect the importance of studying the potential role of microorganisms besides bacteria within microbial mats (Carreira et al., 2020). It is important to mention that metabolomics studies of samples containing diverse communities are scarce; thus, this work significantly expands our understanding of soil metabolites, especially from rare elastic hypersaline microbial mats.

### Conclusions & Perspectives

We found a very low number of taxa that comprised a considerable proportion of the mat and were shared across all sampling points, whereas the rare biosphere was more phylogenetically diverse (FPD index) and phylogenetically disperse (using a null model distribution of Phylogenetic Species Clustering) than the abundant (high read count) taxa for both analyzed libraries. We also found a distinctive metabolome profile for each sample and were able to tentatively annotate several classes of compounds with relevant biological properties.

In view of these results, we are interested to investigate if the metabolomic data is associated with local adaptive processes that could be following an eco-evolutionary dynamic in these fluctuating communities (i.e. taxa are constantly and actively modifying their own ecological niche and that of others (niche construction) (Laland et al., 2016; Odling-Smee et al., 2003, Van Der Hooft et al., 2020)). We also want to explore the high phylogenetic diversity and phylogenetic dispersion shown by the rare biosphere and use transcriptomic data to analyze if the rare biosphere is an active part of this community. We hypothesized that there is a diverse “seed bank” in the aquifer of CCB, a deep biosphere fed by magmatic energy (Wolaver 2013) that could be constantly “seeding” the Archaean Domes community, along with deep water and particular minerals. These ideas could be tested in a deep and spatial-temporal sampling approaches.

## Associated Data

### New DNA/RNA/peptide etc. sequences were reported. Sequences supplied by author here

All raw sequences genaeted are available at GeneBank BioProject PRJNA705156 (BioSample ITS accession SAMN18070760; BioSample 16S accession SAMN18070761). Full data of metabolome feature-based molecular networking can be accessed here: https://massive.ucsd.edu/ProteoSAFe/dataset.jsp?accession=MSV000087545

### Data supplied by the author

Code and processed data are available at GitLab: https://gitlab.com/lauasuar/archaean-domes-repository

## Acknowledgements

We would like to thank Dr. Erika Aguirre-Planter and Jessica Abril Hernández for technical and field assistance. We also thank PRONATURA Noreste for the access to the Pozas Azules ranch, and to Felipe García Oliva (CIEco, Biogeoquímica de suelos) for nutrient analysis of the samples. We acknowledge CONANP (http://sig.conanp.gob.mx/website/pagsig/mapas_serie.htm) and CONABIO SNIB (http://www.conabio.gob.mx/informacion/gis/) public sources utilized to design the Cuatro Ciénegas Basin map in Figure 1.

