## Supplementary material for "Diversity of an uncommon elastic hypersaline microbial mat along a small-scale transect": Suppl. Table 2

List of ITSs ASVs blast hits with <90%

|  | % ITS library |
| --- | --- |
| *Fuscheria nodosa (Alveolata)* | 0.0261 |
| *Enchelys gasterosteus (Alveolata)* | 0.0014 |
| *Strombidium guangdongense (Alveolata)* | 0.0214 |
| *Fabrea salina (Alveolata)* | 0.5524 |
| *Nassula labiata (Alveolata)* | 0.0007 |
| *Paracladotricha salina (Alveolata)* | 0.0207 |
| *Thraustochytrium aureum (Stramenopila)* | 0.0021 |
| *Uncultured Amoebozoa (Amoebozoa)* | 0.0043 |
| *Vannella simplex (Amoebozoa)* | 0.6675 |
| *Chlamydomonadales sp. (Viridiplantae)* | 0.0035 |
| *Hamakko caudatus (Viridiplantae)* | 0.0018 |
| *Eleusine indica (Viridiplantae)* | 0.0078 |
| *Nephelochloa orientalis (Viridiplantae)* | 0.0086 |
