## Supplementary material for "Diversity of an uncommon elastic hypersaline microbial mat along a small-scale transect": Suppl. Table 3

| ITS region kingdom amplicon abundance and ASV number (Forward only database, using UNITE database Eukaryal taxonomy) | S01 | S02 | S03 | S04 | S06 | S07 | | |
|---|---|---|---|---|---|---|---|---|
| Alveolata | 73308 | 53833 | 71131 | 84579 | 155785 | 194989 | | |
| Amoebozoa | 938 | 139 | 433 | 776 | 874 | 2342 | | |
| Protista | 2203 | 3215 | 929 | 458 | 237 | 491 | | |
| Chromista | 0 | 44 | 0 | 0 | 0 | 0 | | |
| Viridiplantae | 25899 | 19608 | 8959 | 5098 | 4150 | 2441 | | |
| Rhodoplantae | 18 | 9 | 26 | 3 | 11 | 22 | | |
| Stramenopila | 0 | 6 | 0 | 0 | 0 | 0 | | |
| Fungi | 6938 | 11471 | 24026 | 11173 | 20383 | 17496 | | |
| Metazoa | 1347 | 2507 | 3147 | 2348 | 1667 | 965 | | |
| TOTAL Classified | 110651 | 90832 | 108651 | 104435 | 183107 | 218746 | 816422 | Sum of classified reads |
| Unclassified | 242069 | 127246 | 59009 | 200803 | 107999 | 71296 | 1624844 | Sum of all reads (ASVs assigned) |
| TOTAL | 352720 | 218078 | 167660 | 305238 | 291106 | 290042 | 50.25% | Percentage classified reads |

| ASV number | | |
|---|---|---|
| Alveolata | 137 | Protozoa ASVs = 329 |
| Amoebozoa | 28 | |
| Protista | 22 | |
| Chromista | 1 | |
| Rhodoplantae | 4 | |
| Stramenopila | 137 | |
| Viridiplantae | 1 | |
| Fungi | 109 | |
| Metazoa | 23 | |
| NA | 461 | |
| TOTAL ASV number | 923 | |

16S rRNA gene phylum amplicon abundance (rarified). Red cells indicate phylum that contains abundant (<1% overall) ASVs

| | | 1 | 2 | 3 | 4 | 5 | 6 | 7 | 8 | 9 | 10 |
|---|---|---|---|---|---|---|---|---|---|---|---|
| ARCHAEA | | | | | | | | | | | |
| 1 | D_0__Archaea;__ | 64 | 60 | 0 | 17 | 11 | 6 | 29 | 8 | 0 | 0 |
| 2 | D_0__Archaea;D_1__Euryarchaeota | 3 | 41 | 0 | 0 | 26 | 97 | 0 | 0 | 0 | 0 |
| 3 | D_0__Archaea;D_1__Woesearchaeota | 8 | 24 | 0 | 27 | 0 | 16 | 60 | 9 | 0 | 6 |
| BACTERIA | | | | | | | | | | | |
| 1 | D_0__Bacteria;D_1__Acetothermia | 129 | 10 | 52 | 15 | 1144 | 2277 | 7 | 0 | 0 | 0 |
| 2 | D_0__Bacteria;D_1__Acidobacteria | 0 | 0 | 0 | 3 | 0 | 0 | 0 | 10 | 31 | 0 |
| 3 | D_0__Bacteria;D_1__Actinobacteria | 8407 | 7669 | 4096 | 6915 | 12210 | 6621 | 4414 | 1849 | 7331 | 3030 |
| 4 | D_0__Bacteria;D_1__Aegiribacteria | 142 | 38 | 0 | 0 | 684 | 10 | 0 | 0 | 0 | 0 |
| 5 | D_0__Bacteria;D_1__Armatimonadetes | 135 | 99 | 120 | 294 | 43 | 420 | 365 | 26 | 126 | 211 |
| 6 | D_0__Bacteria;D_1__Atribacteria | 572 | 153 | 245 | 273 | 7077 | 1322 | 135 | 115 | 13 | 19 |
| 7 | D_0__Bacteria;D_1__Bacteroidetes | 134668 | 153556 | 191232 | 163899 | 125917 | 165242 | 180444 | 281383 | 283266 | 101350 |
| 8 | D_0__Bacteria;D_1__BRC1 | 1777 | 3285 | 2086 | 1602 | 1153 | 1081 | 3394 | 3392 | 231 | 216 |
| 9 | D_0__Bacteria;D_1__Chlamydiae | 281 | 301 | 24 | 122 | 36 | 192 | 379 | 256 | 32 | 0 |
| 10 | D_0__Bacteria;D_1__Chloroflexi | 43304 | 29063 | 6413 | 47211 | 41227 | 44979 | 20777 | 22024 | 41194 | 11244 |
| 11 | D_0__Bacteria;D_1__CK-2C2-2 | 0 | 5 | 0 | 0 | 0 | 0 | 0 | 0 | 0 | 0 |
| 12 | D_0__Bacteria;D_1__Cloacimonetes | 112 | 658 | 58 | 8 | 0 | 0 | 4 | 0 | 0 | 0 |
| 13 | D_0__Bacteria;D_1__Cyanobacteria | 178468 | 118933 | 164747 | 169076 | 77423 | 211897 | 164081 | 129929 | 118366 | 61562 |
| 14 | D_0__Bacteria;D_1__Deinococcus-Thermus | 363 | 496 | 141 | 167 | 277 | 304 | 128 | 218 | 0 | 336 |
| 15 | D_0__Bacteria;D_1__Dependentiae | 13 | 0 | 32 | 62 | 142 | 39 | 96 | 40 | 0 | 0 |
| 16 | D_0__Bacteria;D_1__Elusimicrobia | 32 | 38 | 43 | 45 | 754 | 68 | 34 | 89 | 0 | 0 |
| 17 | D_0__Bacteria;D_1__Epsilonbacteraeota | 6384 | 7925 | 248 | 5249 | 1210 | 1218 | 11976 | 14629 | 2773 | 2836 |
| 18 | D_0__Bacteria;D_1__Fibrobacteres | 4744 | 9465 | 2546 | 7410 | 5276 | 2741 | 3378 | 6510 | 4530 | 1803 |
| 19 | D_0__Bacteria;D_1__Firmicutes | 21350 | 22247 | 15217 | 9825 | 61093 | 12689 | 11840 | 15965 | 15169 | 23660 |
| 20 | D_0__Bacteria;D_1__Fusobacteria | 57 | 33 | 13 | 16 | 20 | 4 | 37 | 23 | 0 | 0 |
| 21 | D_0__Bacteria;D_1__Gemmatimonadetes | 15451 | 13014 | 3766 | 10268 | 10624 | 6286 | 9812 | 9923 | 39397 | 5622 |
| 22 | D_0__Bacteria;D_1__Halanaerobiaeota | 10500 | 17191 | 35438 | 5328 | 40211 | 6557 | 11496 | 30810 | 2480 | 10343 |
| 23 | D_0__Bacteria;D_1__Hydrogenedentes | 351 | 805 | 460 | 400 | 1076 | 325 | 326 | 226 | 13 | 136 |
| 24 | D_0__Bacteria;D_1__Kiritimatiellaeota | 383 | 553 | 271 | 604 | 229 | 315 | 920 | 273 | 17 | 0 |
| 25 | D_0__Bacteria;D_1__Latescibacteria | 755 | 1052 | 219 | 493 | 570 | 557 | 799 | 1215 | 3280 | 1582 |
| 26 | D_0__Bacteria;D_1__LCP-89 | 0 | 0 | 0 | 0 | 0 | 0 | 0 | 6 | 12 | 0 |
| 27 | D_0__Bacteria;D_1__Lentisphaerae | 1972 | 2306 | 2357 | 495 | 1426 | 327 | 1284 | 1457 | 1462 | 257 |
| 28 | D_0__Bacteria;D_1__Margulisbacteria | 166 | 0 | 17 | 26 | 74 | 23 | 100 | 41 | 0 | 3 |
| 29 | D_0__Bacteria;D_1__Marinimicrobia (SAR406 cla | 97 | 906 | 431 | 173 | 283 | 75 | 1657 | 26 | 0 | 0 |
| 30 | D_0__Bacteria;D_1__MAT-CR-M4-B07 | 0 | 70 | 8 | 14 | 0 | 0 | 0 | 0 | 0 | 0 |
| 31 | D_0__Bacteria;D_1__Omnitrophicaeota | 30 | 37 | 74 | 47 | 188 | 23 | 277 | 65 | 0 | 43 |
| 32 | D_0__Bacteria;D_1__Patescibacteria | 17159 | 16264 | 3493 | 31165 | 7895 | 11468 | 28058 | 35446 | 22413 | 20970 |
| 33 | D_0__Bacteria;D_1__Planctomycetes | 12906 | 13262 | 7265 | 14788 | 7614 | 20303 | 16854 | 8469 | 8828 | 2296 |
| 34 | D_0__Bacteria;D_1__Proteobacteria | 169708 | 169915 | 138870 | 157279 | 215777 | 129417 | 152922 | 109274 | 147175 | 59164 |
| 35 | D_0__Bacteria;D_1__Spirochaetes | 77444 | 98384 | 75407 | 79671 | 82623 | 63525 | 63786 | 51066 | 42598 | 10856 |
| 36 | D_0__Bacteria;D_1__Synergistetes | 169 | 350 | 921 | 42 | 1483 | 179 | 80 | 0 | 0 | 0 |
| 37 | D_0__Bacteria;D_1__Tenericutes | 2320 | 7003 | 3500 | 4200 | 3312 | 1124 | 6143 | 10586 | 787 | 3028 |
| 38 | D_0__Bacteria;D_1__Thermotogae | 198 | 3317 | 703 | 455 | 896 | 133 | 1527 | 82 | 0 | 0 |
| 39 | D_0__Bacteria;D_1__Verrucomicrobia | 1524 | 2645 | 614 | 3222 | 468 | 1713 | 2555 | 1140 | 125 | 340 |
| 40 | D_0__Bacteria;D_1__WS1 | 19 | 19 | 0 | 9 | 0 | 21 | 9 | 0 | 0 | 4 |
| UNCLASSIFIED | | | | | | | | | | | |
| 1 | Unassigned;__ | 4980 | 6157 | 996 | 4086 | 227 | 9761 | 7874 | 959 | 1773 | 2587 |
| 2 | D_0__Bacteria;__ | 42855 | 52651 | 97877 | 34999 | 49301 | 56645 | 51943 | 22461 | 16578 | 436496 |
| TOTAL UNCLASSIFIED | | 47899 | 58868 | 98873 | 39102 | 49539 | 66412 | 59846 | 23428 | 18351 | 439083 |
| PERCENTAGE UNCLASSIFIED WITHIN 760000 | | 6.3025 | 7.7457895 | 13.009605 | 5.145 | 6.5182895 | 8.7384211 | 7.8744737 | 3.08263158 | 2.414605263 | 57.77407895 |
| | READS PER SITE (NON RARIFIED) | 945825 | 1164094 | 932904 | 921970 | 1304754 | 850608 | 1056618 | 1444352 | 760326 | 1973982 |

| READS ABUNDANCE | | ASVs NUMBER |
|---|---|---|
| 901401 | Non classified | 4132 |
| 7600000 | Sum of all | 6063 |
| 11.86053947 | Percentage non classified | 68.15 |
