## Supplementary material for "Diversity of an uncommon elastic hypersaline microbial mat along a small-scale transect": Suppl. Table 6

Non-significatives Mantel test results for geographic distance vs beta diversity

16S Bray p=0.0724    r= 0.2749
ITS Jaccard  p= 0.13056   r= 0.3621
ITS Bray  p= 0.13194   r=0.3623

Non-significatives Mantel test results for metabolomic data vs beta diversity

16S Jaccard p= 0.6126 r= -0.08238005
16S Bray p= 0.4274  r= 0.02121974
ITS Jaccard p=0.3611 r=0.0794
ITS Bray p=0.303 r=0.1178
