## Supplementary material for "Diversity of an uncommon elastic hypersaline microbial mat along a small-scale transect": Suppl. Table 7

| | psc_16S | psc_rar_16S | psc_abun_16 | psc_ITS | psc_rar_ITS | psc_abun_ITS |
|---|---|---|---|---|---|---|
| S1 | 0.11060947 | 0.30236487 | 0.04866875 | 0.10968344 | 0.30487551 | 0.10002044 |
| S2 | 0.10863735 | 0.29149669 | 0.0605547 | 0.09490329 | 0.2868876 | 0.10927695 |
| S3 | 0.09063427 | 0.27552425 | 0.05493335 | 0.08496886 | 0.26553793 | 0.12463795 |
| S4 | 0.11408563 | 0.32433312 | 0.05225602 | 0.11582347 | 0.30695845 | 0.09225003 |
| S5 | 0.07809136 | 0.24593114 | 0.04262518 | | | |
| S6 | 0.11513793 | 0.30216016 | 0.08670753 | 0.11553622 | 0.29763445 | 0.1295801 |
| S7 | 0.11590233 | 0.30581557 | 0.07509745 | 0.12709692 | 0.37551424 | 0.11824146 |
| S8 | 0.07955718 | 0.24829185 | 0.04620871 | | | |
| S9 | 0.13062694 | 0.34927144 | 0.07490645 | | | |
| S10 | 0.1354835 | 0.33490847 | 0.08888336 | | | |
