## Supplementary material for "Diversity of an uncommon elastic hypersaline microbial mat along a small-scale transect": Suppl. Table 8

| Suppl. Table 8. Annotated metabolites in samples S1-S10 by feature-based GNPS molecular networking and manual metabolomic analysis. | | | | | |
| --- | --- | --- | --- | --- | --- |
| Observed *m/z* | Exact mass and adduct | Molecular formula | Mass error (in ppm) | Tentative identification (source) | Sample |
| *Microbial metabolites* | | | | | |
| 290.081 | 290.0817 [M+H]^+^ | C_18_H_11_NO_3_ | −0.2 | Nostodione A (*Nostoc commune*) | S2,S6-S8 |
| 318.240 | 318.2433 [M+H]^+^ | C_20_H_31_NO_2_ | +1.7 | Periconiasin G (*Streptomyces* nitrosporeus and *Periconia* sp. F-31) | S1-S3,S6,S7,S9,S10 |
| 319.117 | 319.1182 [M+H]^+^ | C_17_H_18_O_6_ | +1.7 | Dechlorogriseofulvin (several fungal and *Streptomyces* species) | S7 |
| 319.165 | 319.1655 [M+H]^+^ | C_17_H_22_N_2_O_4_ | +0.8 | Phomamide (*Phoma lingam*), Daryamide C (*Streptomyces* spp.), Tomaymycin I (*Nocardia* sp.) | S1-S10 |
| 353.078 | 353.0792 [M+H]^+^ | C_17_H_17_ClO_6_ | +1.6 | Griseofulvin (several fungal and *Streptomyces* species) | S7 |
| 387.180 | 387.1797 [M+H]^+^ | C_22_H_26_O_6_ | −1.3 | Gibbestatin B (*Streptomyces* spp.), Furaquinocin H (*Streptomyces* spp.), Colletofragarone A1 (*Colletotrichum fragariae* and *C. capsiciae*), Pitholide B (*Pestalotiopsis microspore* and *Pithomyces* sp.), Dideacetylparasiticolide A (*Aspergillus oryzae*) | S1-S10 |
| 496.339 | 496.3401 [M+H]^+^ | C_31_H_45_NO_4_ | −4.1 | 8,9-Dihydroindanomycin (*Streptomyces galbus*), Neoindanomycin (*Streptomyces* spp.) | S1-S9 |
| 515.266 | 515.2636 [M+H]^+^ | C_29_H_38_O_8_ | −0.7 | Citreohybridone B (*Penicillium citreoviride*), Setosusin (*Aspergillus duricaulis* and *Corynascus setosus*), Roridin E (several fungal species), Ganoboninone A (*Ganoderma boninense*) | S1-S9 |
| *Plant- and animal-derived metabolites* | | | | | |
| 237.185 | 237.1854 [M+H]^+^ | C_15_H_24_O_2_ | +2.1 | Bisabolene-1,4-endoperoxide (*Commiphora guidottii*) | S1-S3,S6,S7 |
|  | 397.3681 [M+H-H_2_O]^+^ | C_25_H_50_O_4_ | +1.4 | Monobehenin (*Moringa oleifera*) | S5,S9 |
| 353.186 | 353.1865 [M+H]^+^ | C_21_H_24_N_2_O_3_ | +1.5 | Ajmalicine (*Rauvolfia* and *Catharanthus* species) | S7 |
| 233.153 | 233.1541 [M+H]^+^ | C_15_H_20_O_2_ | +2.1 | Costunolide (*Saussurea costus* roots and lettuce) | S1,S3,S9,S10 |
| 204.030 | 204.0297 [M-H]^-^ | C_10_H_7_NO_4_ | -2.7 | Xanthurenic acid (gut of *Anopheles* mosquito) | S1-S9 |
| 311.221 | 311.2225 [M+H-H_2_O]^+^ | C_18_H_32_O_5_ | +2.6 | (10*E*,15*E*)-9,12,13-Trihydroxyoctadeca-10,15-dienoic acid (*Cyperus rotundus*) | S1-S10 |
| 254.284 | 254.2848 [M+H]^+^ | C_17_H_35_N | +2.3 | Solenopsin A (fire ant *Solenopsis invicta*) | S1-S10 |
| 321.242 | 321.2429 [M+H]^+^ | C_20_H_32_O_3_ | +1.7 | 6-Oxocativic acid (*Cistus ladaniferus*) | S1-S10 |
| 295.227 | 295.2273 [M+H]^+^ | C_18_H_30_O_3_ | +1.8 | 13-Keto-9*Z*,11*E*-octadecadienoic acid (*Carthamus oxyacantha*) | S2,S4-S9 |
| *Human activity-derived chemicals* | | | | | |
| 377.214 | 377.2148 [M+H]^+^ | C_14_H_29_N_6_O_6_ | +1.3 | Hexamethylolmelamine pentamethyl ether (water contaminant) | S2,S4,S6-S8,S10 |
| 288.253 | 288.2539 [M+H]^+^ | C_16_H_33_NO_3_ | +1.9 | Lauryl diethanolamide (cosmetics) | S8,S9 |
| 284.295 | 284.2954 [M+H]^+^ | C_18_H_37_NO | +2.1 | Octadecanamide (surfactant and emulsifier) | S3,S8 |
