## Supplementary material for "Diversity of an uncommon elastic hypersaline microbial mat along a small-scale transect": Suppl. Table 10

Acinetobacter (Proteobacteria) 1
Aerococcus (Firmicutes)
Albimonas (Proteobacteria)
Aliifodinibius (Bacteroidetes)
Alkaliflexus (Bacteroidetes)
Allofrancisella (Proteobacteria)
Anaerobranca (Firmicutes)
Anaerosalibacter (Firmicutes)
Aquicella (Proteobacteria)
Bacillus (Firmicutes)
bacterium YC-ZSS-LKJ186 (Aegiribacteria)
Bdellovibrio (Proteobacteria)
Blastopirellula (Planctomycetes)
Brevinema (Spirochaetes)
Brochothrix (Firmicutes)
Candidatus (Paenicardinium (Bacteroidetes)
Candidatus Endoecteinascidia (Proteobacteria)
Candidatus Endomicrobium (Elusimicrobia)
Candidatus Fritschea (Chlamydiae)
Candidatus Megaira (Proteobacteria)
Candidatus Melainabacteria bacterium RIFOXYA2_FULL_32_9 (Cyanobacteria)
Candidatus Phytoplasma (Tenericutes)
Carboxydocella (Firmicutes)
Cerasicoccus (Verrucomicrobia)
Chloroflexi bacterium RBG_13_46_14 (Chloroflexi)
Clostridiales bacterium PH28_bin88 (Chloroflexi)
Clostridiisalibacter (Firmicutes)
Clostridium sensu stricto 7 (Firmicutes)
Defluviitalea (Firmicutes)
Defluviitaleaceae UCG-011 (Firmicutes)
Dehalobium (Chloroflexi)
Desulfocella (Proteobacteria)
Desulfonatronovibrio (Proteobacteria)
Desulfuribacillus (Firmicutes)
Enterococcus (Firmicutes)
Escherichia coli (Proteobacteria)
Estrella (Chlamydiae)
FFCH7168 (Chloroflexi)
Fodinicurvata (Proteobacteria)
Francisella (Proteobacteria)
Garciella (Firmicutes)
Geitlerinema PCC-7105 (Cyanobacteria) 2
Halarsenatibacter (Halanaerobiaeota)
Haliea (Proteobacteria)
Halonatronum (Halanaerobiaeota)
Holospora (Proteobacteria)
Hydrogenispora (Firmicutes)
Lachnospiraceae NK3A20 group (Firmicutes)
Legionella (Proteobacteria)
Leptococcus JA-3-3Ab (Cyanobacteria)
Leuconostoc (Firmicutes)
Lewinella (Bacteroidetes)
Litoricola (Proteobacteria)
Lutispora (Firmicutes)
MAT-CR-P4-C12 (Bacteroidetes)
Melioribacter (Bacteroidetes)
Modestobacter (Actinobacteria)
MSBL3 (Kiritimatiellaeota)
Neochlamydia (Chlamydiae)
Nitrolancea (Chloroflexi)
Ohtaekwangia (Bacteroidetes)
Oscillatoria PCC-6304 (Cyanobacteria)
Paenibacillus (Firmicutes)
Parcubacteria group bacterium GW2011_GWF2_45_11 (Patescibacteria)
Phaselicystis (Proteobacteria)
Planktosalinus (Bacteroidetes)
Prevotella 1 (Bacteroidetes)
Prevotella 7 (Bacteroidetes)
Prevotellaceae UCG-003 (Bacteroidetes)
Pseudobacteriovorax (Proteobacteria)
Pseudobacteroides (Firmicutes)
Pseudohongiella (Proteobacteria)
Pseudomonas (Proteobacteria)
Psychrobacter (Proteobacteria)
Psychroflexus (Bacteroidetes)
Rhodococcus (Actinobacteria)
Rhodopirellula (Planctomycetes)
Rikenellaceae RC9 gut group (Bacteroidetes)
Roseiflexus (Chloroflexi)
Roseimarinus (Bacteroidetes)
Roseobacter clade CHAB-I-5 lineage (Proteobacteria)
Rubellimicrobium (Proteobacteria)
Rubrobacter (Actinobacteria)
Ruminiclostridium 1 (Firmicutes) 3
Ruminococcaceae NK4A214 group (Firmicutes)
Ruminococcaceae UCG-013 (Firmicutes)
Ruminococcaceae UCG-014 (Firmicutes)
Salinisphaera (Proteobacteria)
Salipaludibacillus (Firmicutes)
Salisediminibacterium (Firmicutes)
SEEP-SRB1 (Proteobacteria)
Silvanigrella (Proteobacteria)
ST-3K26 (Bacteroidetes)
Streptomyces (Actinobacteria)
Sulfurimonas (Epsilonbacteraeota)
Taeseokella (Bacteroidetes)
Thalassospira (Proteobacteria)
Thermovirga (Synergistetes)
Thermus (Deinococcus-Thermus)
Tistrella (Proteobacteria)
Veillonella (Firmicutes)
Wenzhouxiangella (Proteobacteria)
Wohlfahrtiimonas (Proteobacteria)
Geotoga (Thermotogae)
bacterium enrichment culture clone E23 (Cloacimonetes)
Marinococcus (Firmicutes)
Acholeplasma (Tenericutes)
Salinispira (Spirochaetes)
Salinivibrio (Proteobacteria)
Chitinivibrio (Fibrobacteres)
F1-37X2 (Bacteroidetes)
Pontibacter (Bacteroidetes)
ESFC-1 (Cyanobacteria)
Fictibacillus (Firmicutes)
Anaerolinea (Chloroflexi)
Fusibacter (Firmicutes)
Caenispirillum (Proteobacteria)
Sodalis (Proteobacteria)
Halofilum (Proteobacteria)
Desulfofustis (Proteobacteria)
Puniceicoccus (Verrucomicrobia)
Alkaliphilus (Firmicutes)
Rubrivirga (Bacteroidetes)
Candidatus Melainabacteria bacterium GWA2_34_9 (Cyanobacteria)
Desulfotomaculum (Firmicutes)
Oscillochloris (Chloroflexi) 4
Halochromatium (Proteobacteria)
Thiohalorhabdus (Proteobacteria)
Cesiribacter (Bacteroidetes)
Izimaplasma (Tenericutes)
IheB3-7 (Bacteroidetes)
Brachyspira (Spirochaetes)
Pseudokineococcus (Actinobacteria)
bacterium YC-ZSS-LKJ145 (Chloroflexi)
SBZC-1223 (Lentisphaerae)
Peredibacter (Proteobacteria)
Desulfonauticus (Proteobacteria)
Marinobacter (Proteobacteria)
Dehalobacter (Firmicutes)
GWE2-31-10 (Spirochaetes)
Marivita (Proteobacteria)
Ruminiclostridium (Firmicutes)
Chloronema (Chloroflexi)
Candidatus Alysiosphaera (Proteobacteria)
Desulfotignum (Proteobacteria)
Longimonas (Bacteroidetes)
Phaeodactylibacter (Bacteroidetes)
Methylobacterium (Proteobacteria)
Candidatus Latescibacter (Latescibacteria)
Salinibacter (Bacteroidetes)
Owenweeksia (Bacteroidetes)
Phormidium MBIC10003 (Cyanobacteria)
Halopeptonella (Proteobacteria)
Natronincola (Firmicutes)
Desulfatiglans (Proteobacteria)
Parvularcula (Proteobacteria)
Rapidithrix (Bacteroidetes)
Deltaproteobacteria bacterium RIFOXYB12_FULL_58_9 (Proteobacteria)
Pelagicoccus (Verrucomicrobia)
Microgenomates group bacterium RIFCSPLOWO2_01_FULL_47_10 (Patescibacteria)
Desulfonatronum (Proteobacteria)
Lentimonas (Verrucomicrobia)
Orenia (Halanaerobiaeota)
Thiomicrospira (Proteobacteria)
Alterococcus (Verrucomicrobia)
Roseibaca (Proteobacteria)
Caminicella (Firmicutes)
Dethiosulfatibacter (Firmicutes) 5
SM1A02 (Planctomycetes)
Natranaerobius (Firmicutes)
Halorhodospira (Proteobacteria)
Peptoniphilus (Firmicutes)
Halocella (Halanaerobiaeota)
Desulfosalsimonas (Proteobacteria)
Truepera (Deinococcus-Thermus)
Dethiobacter (Firmicutes)
Ruminococcaceae UCG-012 (Firmicutes)
Haloplasma (Tenericutes)
SC103 (Thermotogae)
Planctomycetales bacterium Ellin7244 (Planctomycetes)
Halothiobacillus (Proteobacteria)
Gracilimonas (Bacteroidetes)
[Cytophaga] xylanolytica group (Bacteroidetes)
Alkalibacterium (Firmicutes)
Rhodovulum (Proteobacteria)
Cryomorpha (Bacteroidetes)
Candidatus (Amoebophilus (Bacteroidetes)
Verruc-01 (Verrucomicrobia)
Urania-1B-19 marine sediment group (Planctomycetes)
Rubribacterium (Proteobacteria)
Algoriphagus (Bacteroidetes)
Desulfitibacter (Firmicutes)
bacterium YC-ZSS-LKJ74 (Chloroflexi)
Roseivivax (Proteobacteria)
CENA518 (Cyanobacteria)
Sandaracinus (Proteobacteria)
Flavobacterium (Bacteroidetes)
ADurb.Bin120 (Chloroflexi)
OM27 clade (Proteobacteria)
Candidatus Chlorothrix (Chloroflexi)
Desulfovibrio (Proteobacteria)
Algisphaera (Planctomycetes)
Halomonas (Proteobacteria)
ML310M-34 (Bacteroidetes)
Oceanicaulis (Proteobacteria)
Phycisphaera (Planctomycetes)
Blvii28 wastewater-sludge group (Bacteroidetes)
Rhodohalobacter (Bacteroidetes)
Dactylococcopsis PCC-8305 (Cyanobacteria)
Alkalispirochaeta (Spirochaetes)
