## Supplementary figures and images for "Diversity of an uncommon elastic hypersaline microbial mat along a small-scale transect"

### Suppl Fig 3A1

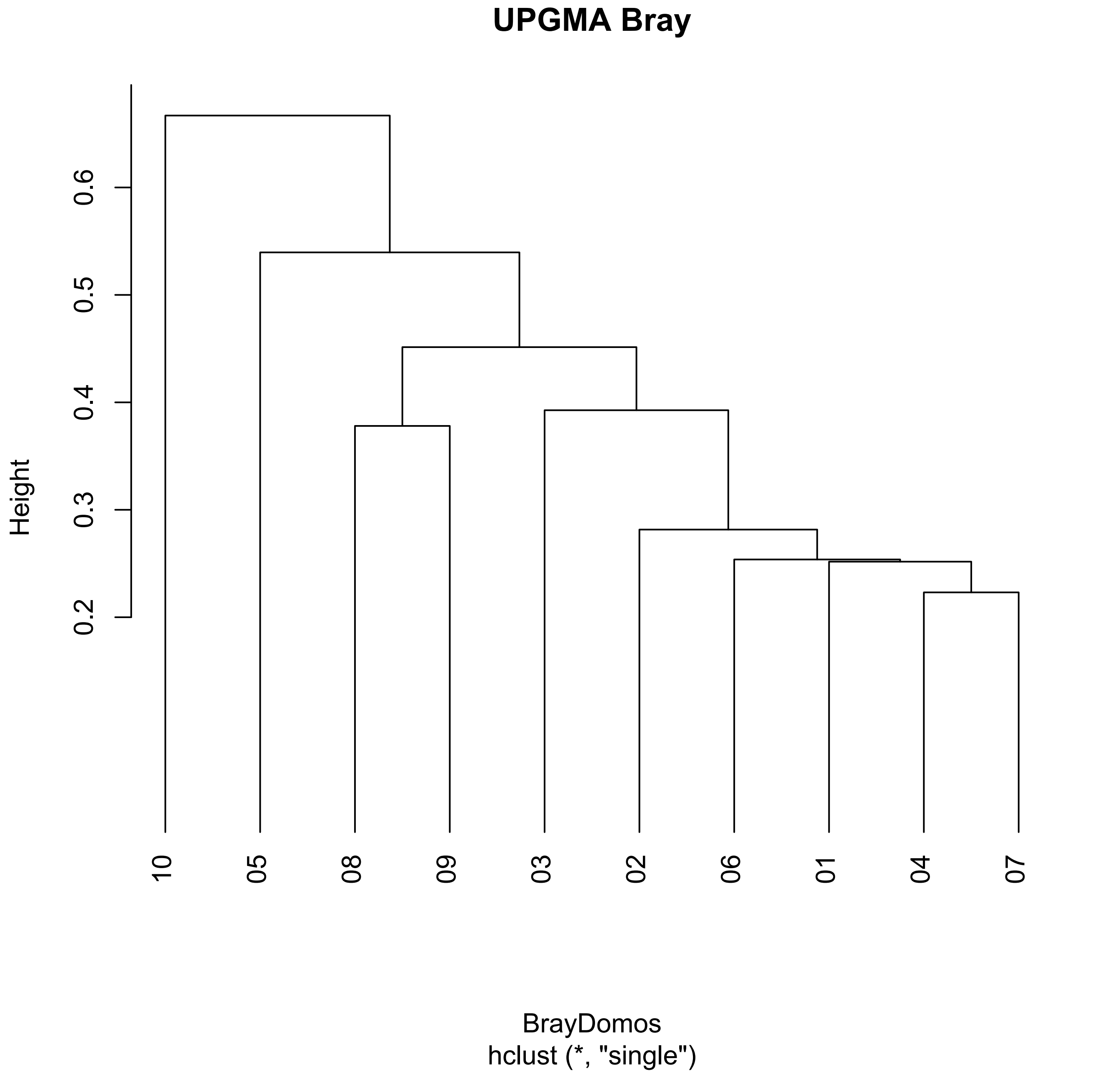

### Suppl Fig 3A2

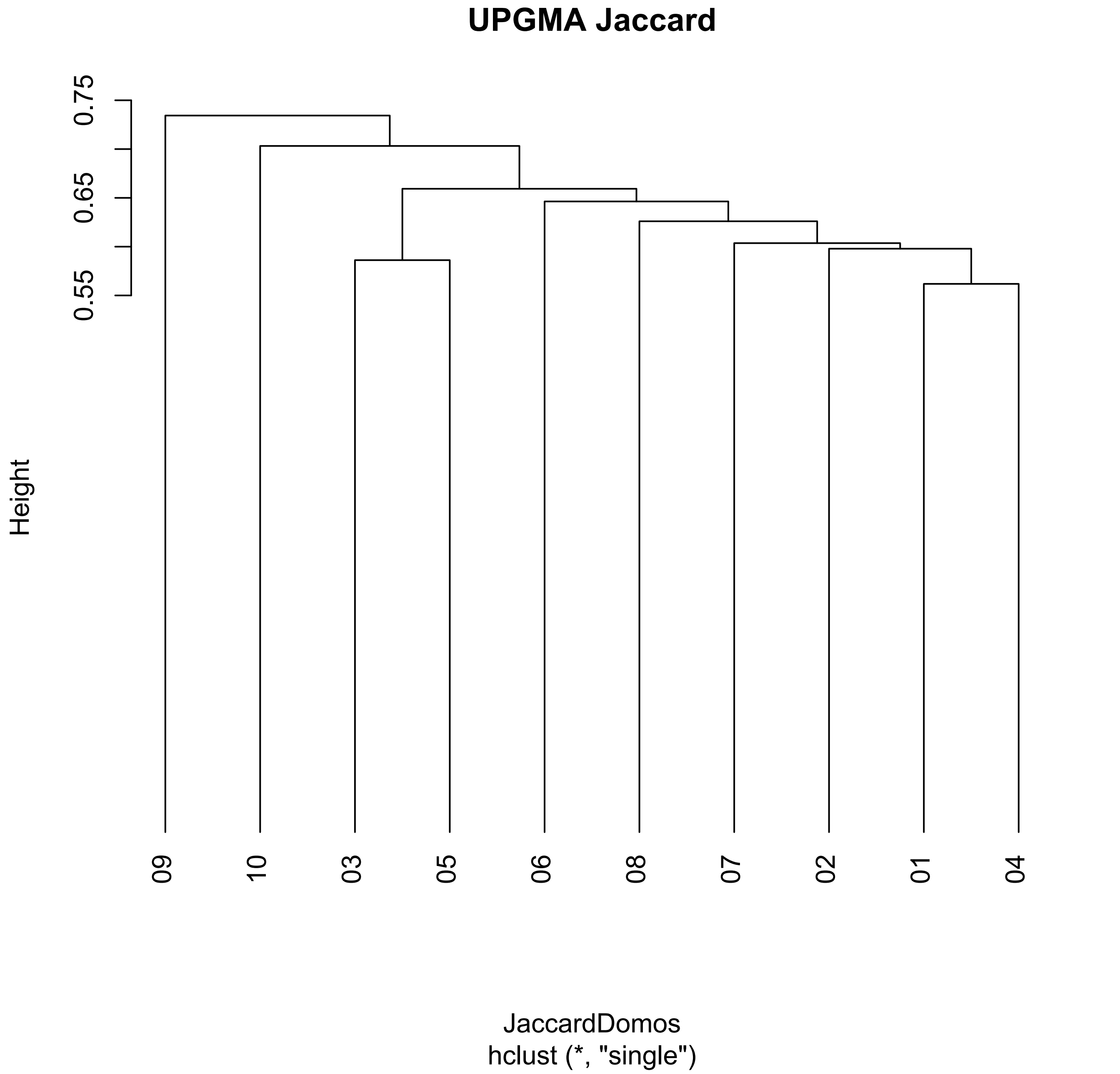

### Suppl Fig 3B1

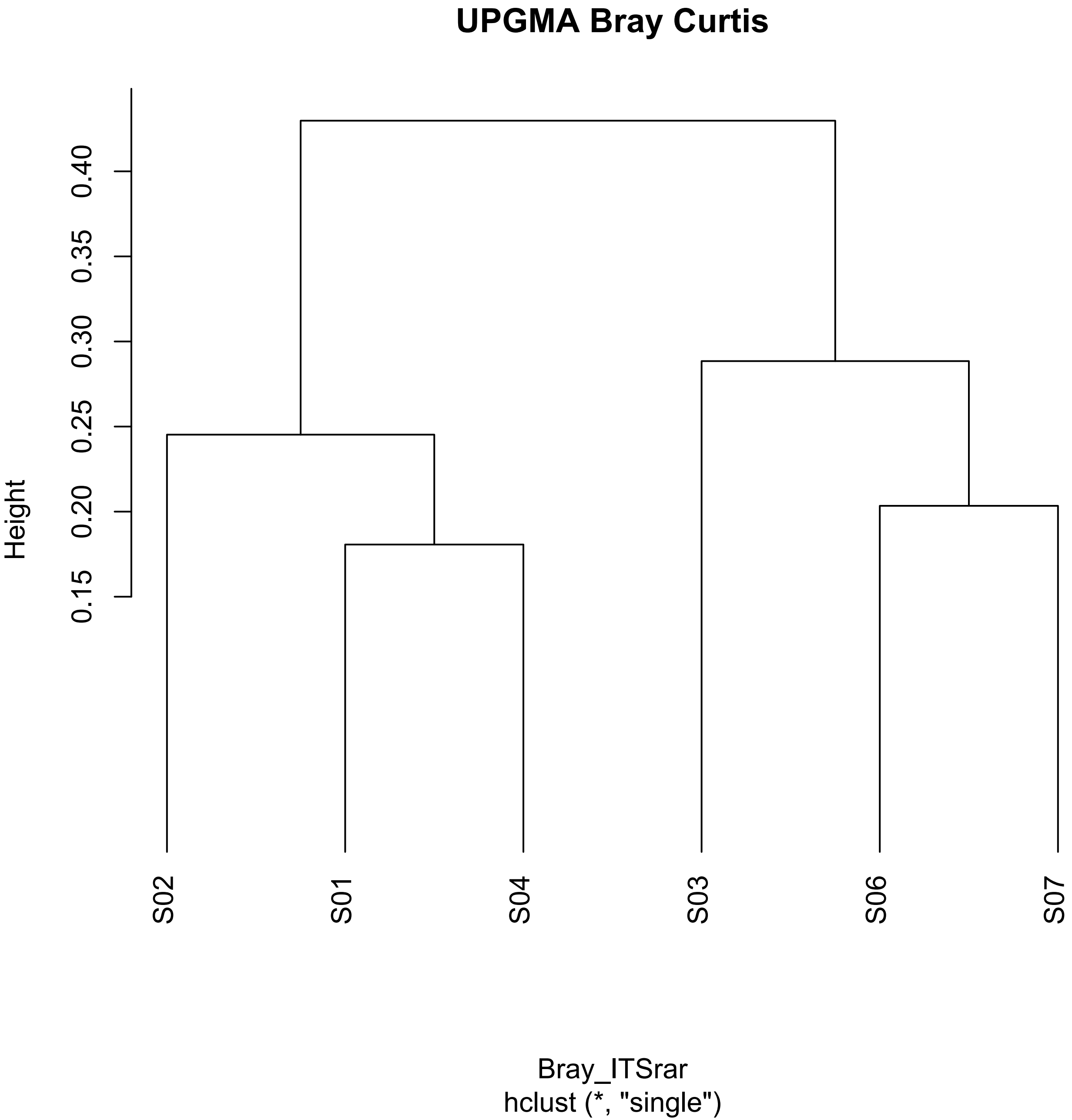

### Suppl Fig 3B2

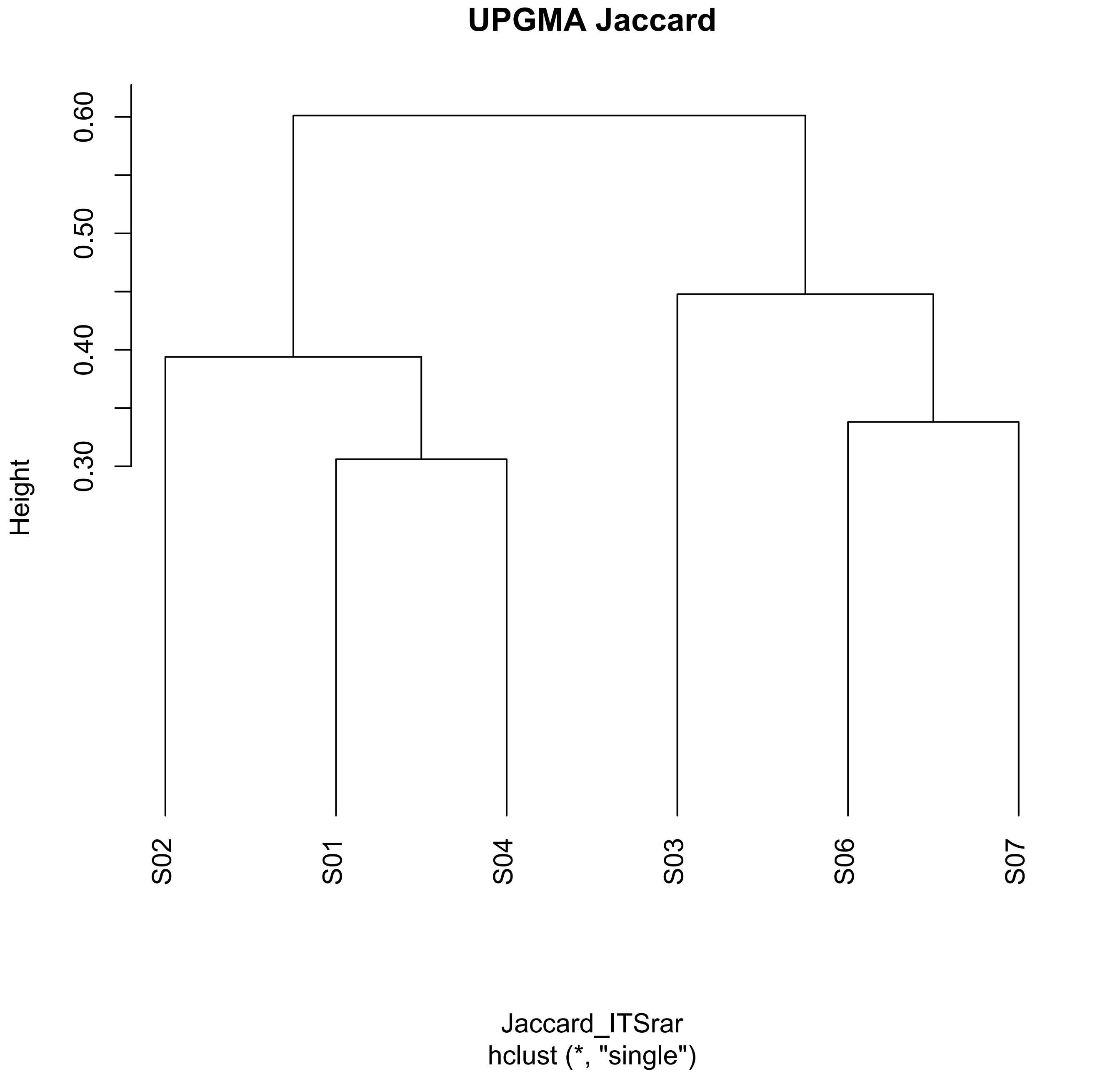

### Suppl Fig 4

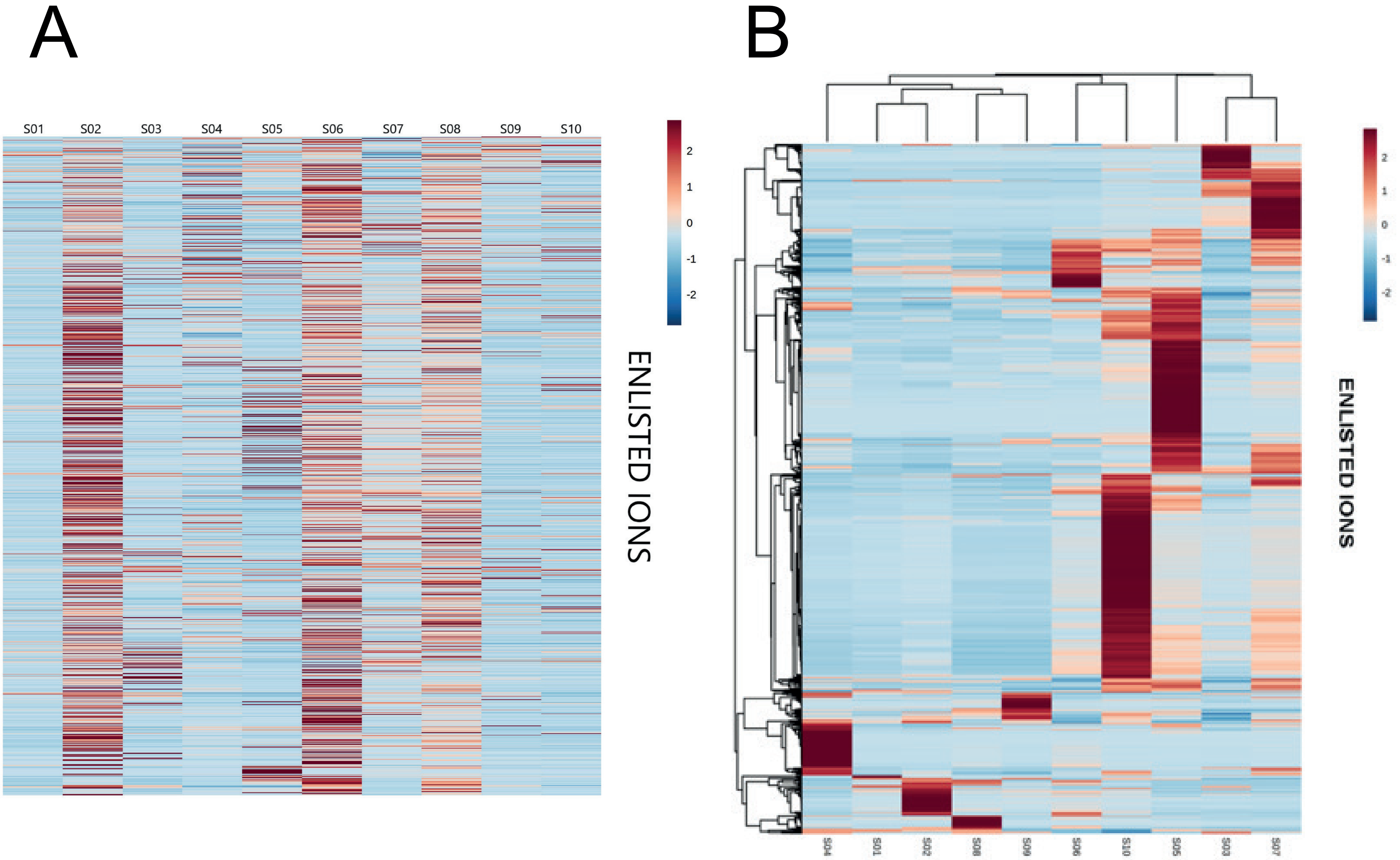

### Suppl. Fig 1A

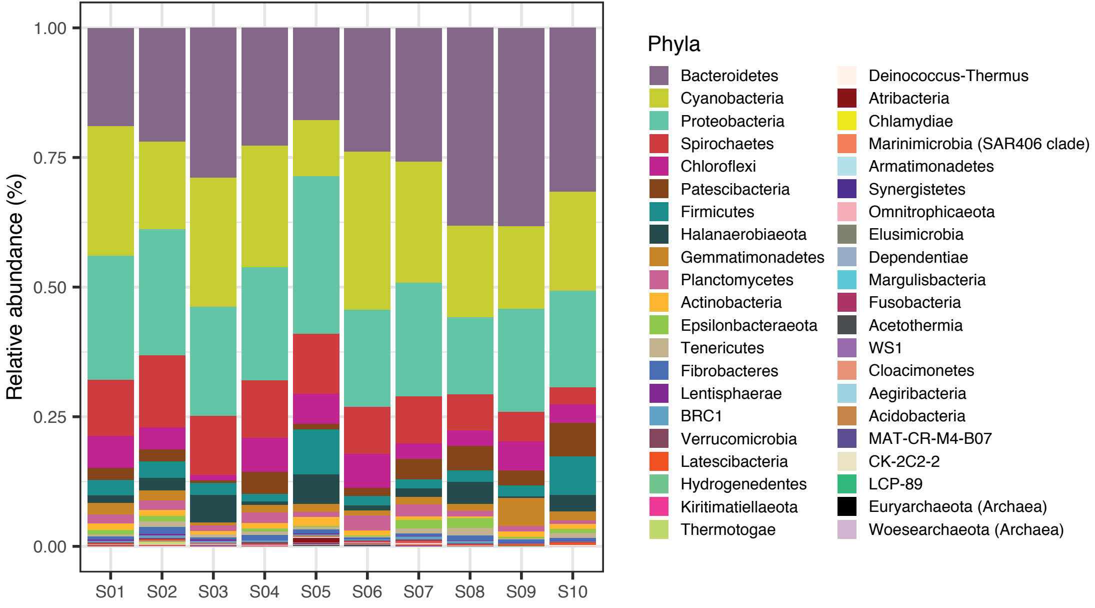

### Suppl. Fig 1B

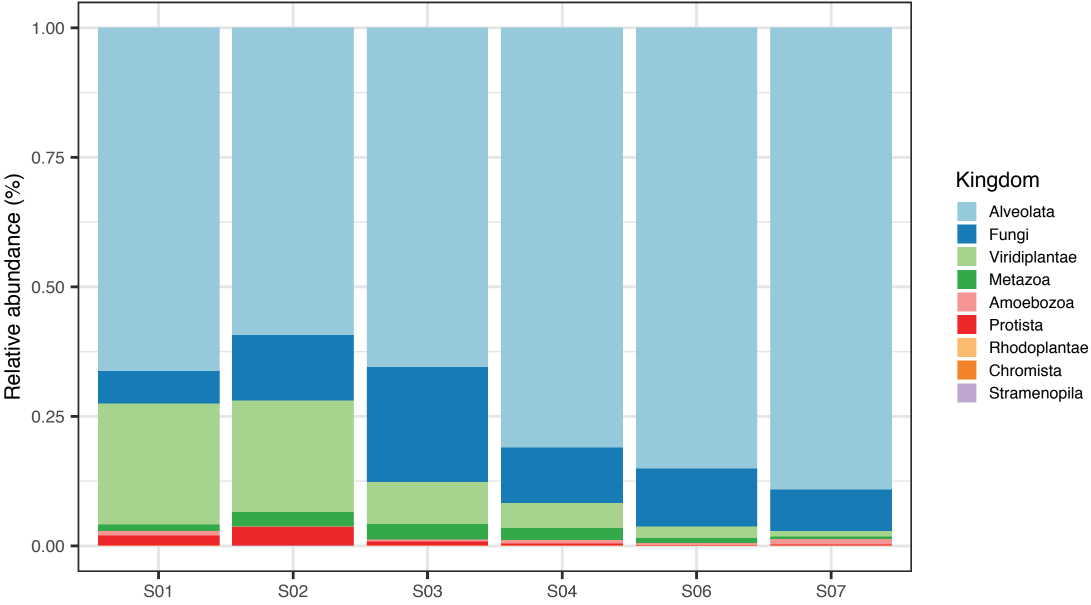

### Suppl. Fig 2A

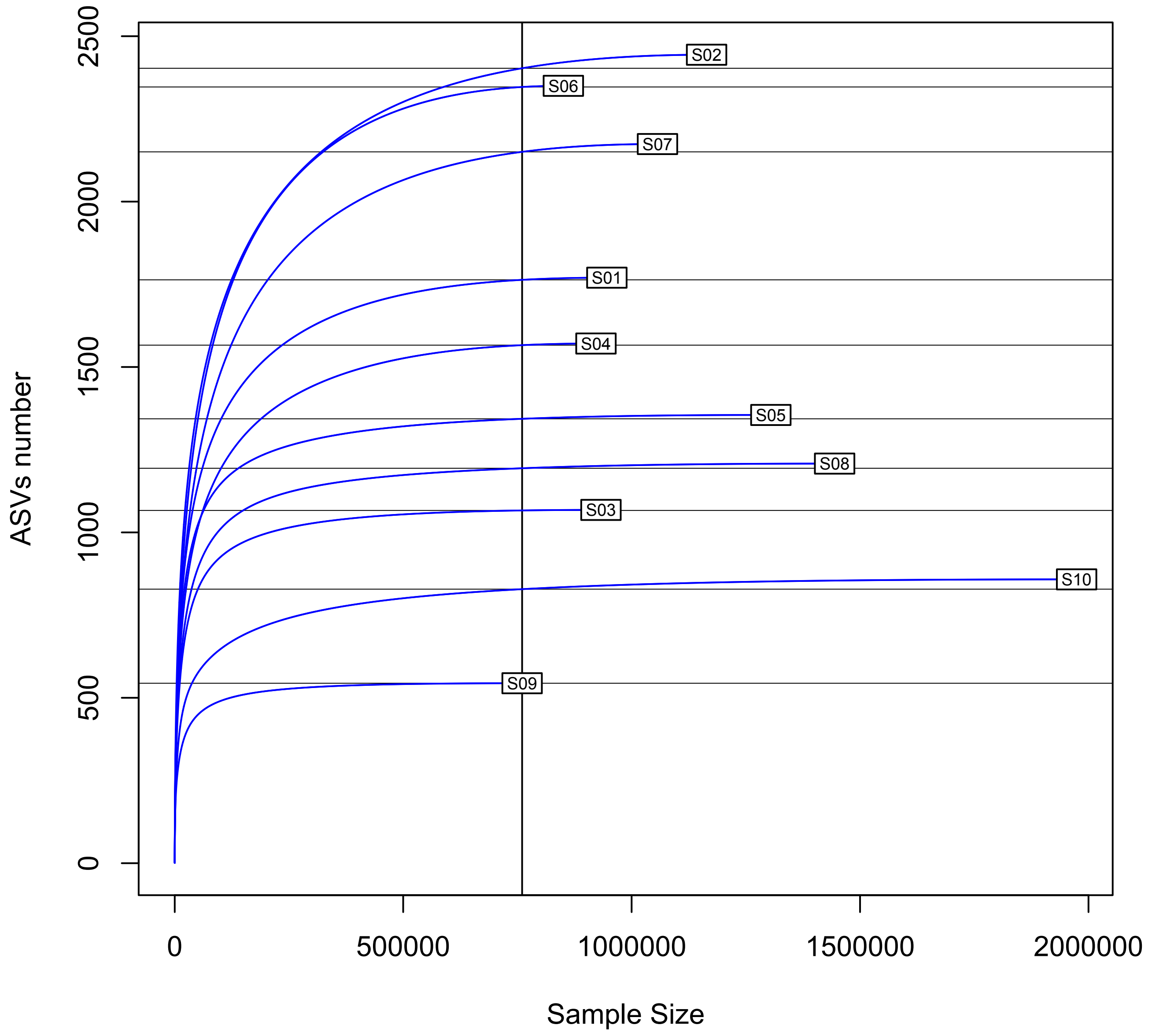

### Suppl. Fig 2B

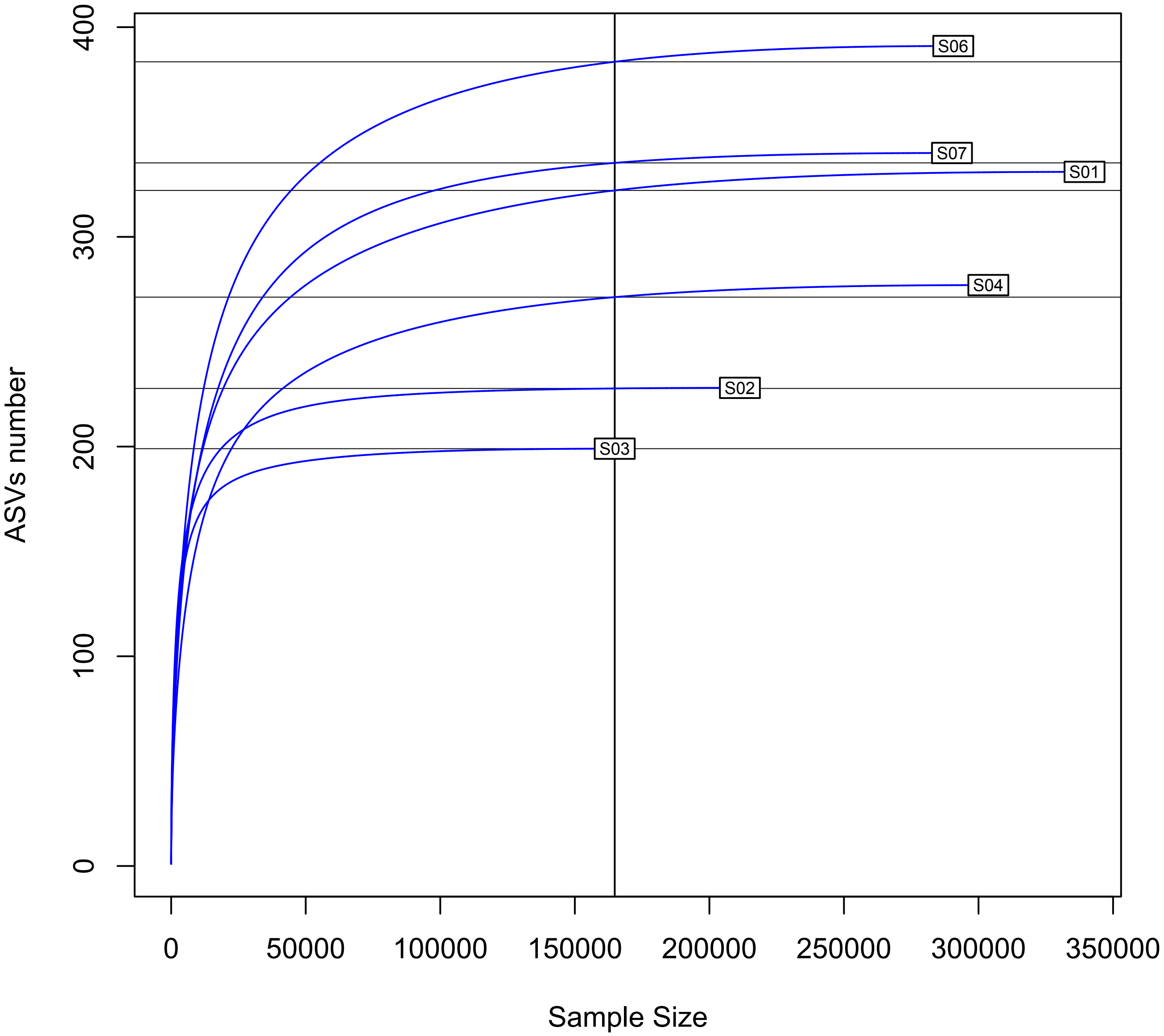
