## Supplementary material for "Diversity of an uncommon elastic hypersaline microbial mat along a small-scale transect": Figure legends

**Table1**

Alpha diversity indices for 16S rDNA data and Forward-Only ITS region data from Archaean Domes CCB microbial mats

**Suppl. Fig. 1**

Taxa relative proportion.

(A) Phylum composition within 16S rDNA gene data (B) Kingdom composition (based on UNITE database eukaryotic taxonomy) within Forward-Only ITS region data.

**Suppl. Fig. 2**

Rarefaction curves.

16S rDNA gene data (A) and ITS region Forward-Only ITS data (B).

**Suppl. Fig. 3**

UPGMA dendrogram showing composition similarities among sampling sites.

Branch length represents Bray-Curtis and Jaccard-1 dissimilarity coefficient, based on (A) 16S rDNA ASVs and (B) Forward-Only ITS ASVs

**Suppl. Figure 4**

Metabolite heatmap analysis.

Relative proportion of metabolite features (MS ions in positive mode) in the Archaean Domes mats samples. On the scale bar, brick red color indicates increased metabolite levels, and blue color represents decreased levels.

(A) non-clustered samples and sites, (B) clustered samples and sites.

**Suppl. Table 1**

Merge ITS-ASV community composition and taxonomy table

**Suppl. Table 2**

List of ASVs Blast hits with <90%, of non-fungal Merge ITS data.

**Suppl. Table 3**

Phyla and kingdom community composition tables

16S rDNA gene phyla and ITS region kingdom taxa community composition ASVs tables, and percentage of classified and unclassified ASVs.

**Suppl. Table 4**

Forward-Only ITS-ASV community composition and taxonomy table

**Suppl. Table 5**

16S rDNA-ASV community composition and taxonomy table

**Suppl. Table 6**

Mantel test negative results of metabolites distance matrix vs beta diversity distance matrix.

**Suppl. Table 7**

nmdPSC values

**Suppl. Table 8**

Identified microbial metabolites.

Tentatively identified microbial metabolites in samples S1-S10 by GNPS and detailed metabolomic analysis.

**Suppl. Table 9**

List of 18 orders shared between Archaean Domes and other layered microbial communities

**Suppl. Table 10**

Labels of Figure 2A (up).

Low read count genera (less than 1% of total proportion of taxa) labels from Figure top 2A. Five repeated sequences of colors are used, distinguished here by red lines, and numerated red arrows in figure.
